# K-Ras controls asymmetric cell divisions from the primary cilium

**DOI:** 10.64898/2026.03.05.709789

**Authors:** Rohan Chippalkatti, Elisabeth Schaffner-Reckinger, Anthoula Gaigneaux, Bianca Parisi, Sara Bottone, Christina Laurini, Yashar Rouzbahani, Mariska Dijkers, Atanasio Gómez-Mulas, Thomas Sauter, Jeroen den Hertog, Christian Eggeling, Daniel Kwaku Abankwa

## Abstract

The Ras-MAPK pathway drives central cellular processes, including cell proliferation and differentiation. How exactly Ras controls differentiation is however not understood. Supported by mathematical modelling and single-cell RNA sequencing we show that K-Ras4B sustains ciliation during differentiation thus restricting commitment of skeletal muscle stem and progenitor cells during asymmetric cell divisions. Modulation of K-Ras4B abundance or expression of oncogenic K-Ras4B-G12C perturb normal differentiation. K-Ras4B, but not N-Ras and H-Ras, localizes to the primary cilium and its abundance there depends on the ciliary trafficking chaperone PDE6D. The presence of B-Raf and active MEK at the base of and active ERK inside the cilium suggests that K-Ras4B is active there. Conditions that localize a K-Ras4B mutant only to the cilium are sufficient to sustain ciliation and normal differentiation. Finally, in vivo modulation of K-Ras4B activity during zebrafish embryogenesis perturbs ciliation-dependent heart-looping. Our results thus imply a novel fundamental role of K-Ras4B in controlling ciliation and differentiation and suggest an explanation for the phenotypic similarities between RASopathies and ciliopathies.

## Introduction

It is currently not fully understood how the Ras pathway controls cell differentiation, a process essential for organismal development and intertwined with the cell cycle ^1^. The profound impact of Ras on human development becomes obvious in RASopathies, rare developmental diseases caused by Ras pathway mutations that result in cardiac, neuronal and musculo-skeletal defects ^2^.

Stem and progenitor cells sit at the apex of differentiation hierarchies. Asymmetric cell divisions of stem and progenitor cells are crucial for tissue maintenance, as this process produces one daughter stem cell to maintain the stem cell pool and one daughter cell committed to differentiate further. Asymmetric cell division requires asymmetric segregation of molecular determinants, such as polarity proteins or fate determinants. Similarly, several cellular organelles, such as the mother centrosome, have been identified to segregate with the cell that retains stem cell properties or stemness traits ^3^. The molecular factors of the mother centrosome that impart stemness traits are currently unknown.

Skeletal muscle provides a well-studied model of cell differentiation. The tissue-resident stem cells in vertebrate muscles are called satellite cells, which are positive for the lineage-specific transcription factor Pax7. Muscle-specific transcription factors, such as Myf5, Myod1 and Myog become sequentially activated to drive terminal differentiation of muscle cells into fused muscle fibers, which can be detected by myosin heavy chain (MyHC) expression ^4^. Asymmetric divisions of satellite cells in vivo generate Pax7+/Myf5-stem cells and Pax7+/Myf5+ progenitors committed to differentiation ^5^. In vivo, primary cilia mark muscle stem cells for self-renewal ^6^, suggesting a role in their asymmetric cell division.

The primary cilium is an antenna-like organelle on stem and progenitor cells. It emerges from the basal body, which is directly derived from the mother centriole of the mother centrosome. Major stemness and developmental pathways such as Hedgehog, Wnt and Notch localize only to the cilium ^7^. During stem and progenitor cell divisions the mother centrosome can remain associated with the membrane remnant of the cilium, which facilitates ciliogenesis in the inheriting daughter cell and sensitizes it for stemness signaling thus imparting stemness traits in an asymmetric fashion ^8^. Therefore, molecular processes that are consolidated in the primary cilium may be instructive for re-ciliation, thus protecting stemness traits during asymmetric divisions of stem cells ^3^.

A spectrum of rare diseases, called ciliopathies, is associated with ciliary dysfunction and display developmental defects overlapping with those seen in RASopathies, including musculo-skeletal and neuro-cognitive abnormalities ^9, 10^. However, a common mechanism that would explain these shared phenotypes between RASopathies and ciliopathies is not known. Many ciliopathy genes regulate ciliary access of proteins. A ciliary gate maintains a diffusion barrier between the cilium and the rest of the cell, allowing passive diffusion of proteins <70 kDa into the cilium ^11^. Lipid-modified proteins require trafficking chaperones UNC119A/B and PDE6D, which release cargo via allosteric release factors Arl2/3 ^12, 13^.

GTP-bound Arl2 can only eject cargo with lower, micromolar or submicromolar affinities to PDE6D, such as Src- and Ras-proteins ^12, 13^. Binding affinity depends on residues near the prenylated cysteine, explaining for instance why K-Ras4A is not a cargo of PDE6D and reversible palmitoylation of N-Ras and H-Ras makes them worse cargos than K-Ras4B ^14, 15,16^. GTP-Arl2 in the perinuclear area releases PDE6D-bound K-Ras4B, which is subsequently trapped on the recycling endosomes for vesicular forward trafficking to the plasma membrane. However, plasma membrane delivery of K-Ras4B appears to occur via redundant pathways as only 25-50 % are performed by PDE6D ^17, 18^. The guanine nucleotide exchange factor (GEF) of related Arl3 is the ciliary marker protein Arl13B, a G domain-containing protein that is likewise active only if GTP-bound ^19^. Arl13B is anchored to the ciliary membrane but also contacts the microtubule core of the cilium ^7^. For both UNC119B and PDE6D, nanomolar, high-affinity cargo such as NPHP3 and INPP5E, respectively, can only be offloaded by GTP-Arl3. However, also low affinity cargo such as the Ras-related small GTPase Rheb is released by Arl3 from PDE6D ^20^.

Supported by mathematical modelling, we here provide evidence that Ras proteins have an isoform-specific impact on ciliation, wherein K-Ras4B regulates asymmetric cell divisions during skeletal muscle cell differentiation. Single-cell RNA sequencing data suggest that a high *KRAS* expression is required for re-ciliation of proliferating stem and progenitor cells. However, both decreased K-Ras4B levels and oncogenic K-Ras4B disrupt normal differentiation, supporting that Ras signaling needs to follow a discrete regulation during differentiation. Aided by the trafficking chaperone PDE6D, K-Ras4B can freely diffuse into the cilium and is active there as supported by the localization of active MAPK-components at the cilium. A mutant that only binds to the membrane of the cilium is sufficient to sustain normal ciliation and differentiation. Expression of dominant negative K-Ras4B-S17N in zebrafish embryos leads to heart-looping defects, which are associated with reduced ciliogenesis in the left-right organizer. Our data suggest that K-Ras4B sustains ciliation and thus restricts commitment during asymmetric cell divisions of skeletal muscle cells. The potentially broad conservation of this mechanism in vertebrate development provides an explanation for the phenotypic similarities between RASopathies and ciliopathies.

## Results

### Mathematical modelling and experimental validation suggest that K-Ras protects muscle stem cells by sustaining their re-ciliation during asymmetric cell divisions

To investigate the ciliary functions of Ras in the context of cell differentiation, we employed the heterogenous murine C2C12 skeletal muscle cell line, which contains a fraction of ciliated cells and can be induced to differentiate into myocytes and muscle fibers within 3-5 days in low serum medium (**Figure 1A, Figure S1A**). This cell line is easy to manipulate chemo-genetically and is uniquely suited to correlate subcellular localization studies and cell differentiation processes that closely align with those seen in vivo ^21^.

**Figure 1.**
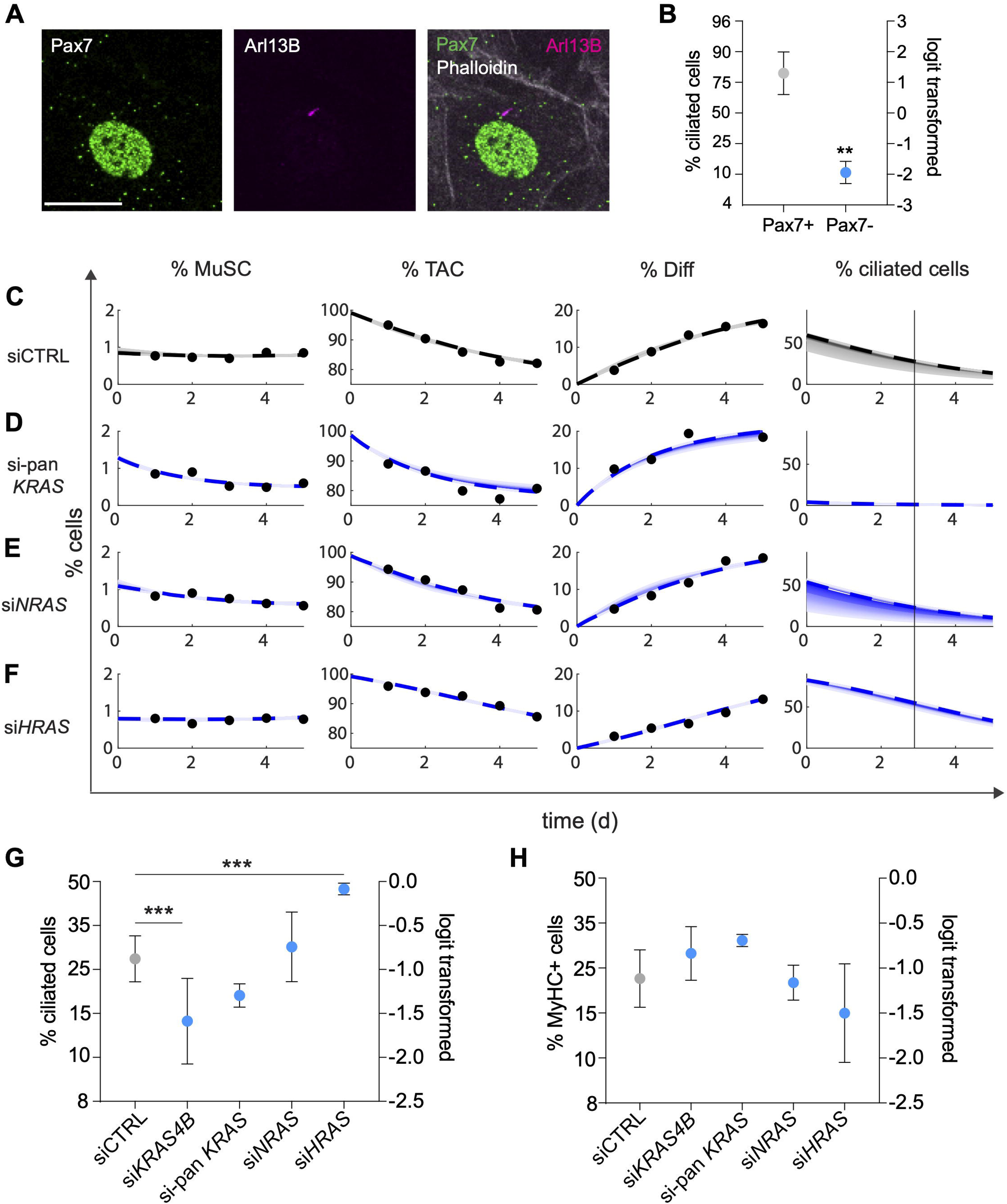
Mathematical modelling and its experimental validation support that K-Ras prevents commitment and differentiation by sustaining primary cilia. **(A)** Confocal images of C2C12 cells grown in high serum for 72 h and immunolabelled for Pax7 in the nucleus and ciliary marker Arl13B. Phalloidin staining visualizes cell outlines. Scale bar = 20 µm. **(B)** Quantification of ciliation using data as in (A), N = 3 independent biological repeats with a total of n = 800 cells. Means ± SD are plotted. Statistical analysis was done with the unpaired t-test. (**C-F**) Modelled (curves) and measured (points) of the population evolution of MuSC, TAC and differentiated C2C12 cells (Diff) after serum switching at day 0 to trigger differentiation. Experimental samples were treated at day -1 with non-targeting siRNA (siCTRL, C), siRNA targeting *KRAS4A/4B* (si-pan KRAS, D), *NRAS* (E) or *HRAS* (F). Plots to the right show predicted percentage of ciliated cells over time with the line indicating day 3 in differentiating low serum. **(G)** Confocal imaging-based quantification of ciliation of C2C12 transfected with the indicated siRNA (100 nM) and grown in low serum for 72 h. N = 3, n > 500. Means ± SD are plotted. Statistical analysis was done with one-way ANOVA and the Dunnett’s T3 multiple comparisons test. **(H)** Flow cytometric quantification of MyHC terminal differentiation marker expression in C2C12 treated as in (G), N = 3. Means ± SD are plotted. Statistical analysis was done with one-way ANOVA and Dunnett’s T3 multiple comparisons test.

We recently provided evidence that the C2C12 skeletal muscle cell line is heterogenous and contains 1-5 % mitotically highly active Pax7+/ MyHC-muscle stem cells (**MuSC**), which under high serum conditions divide asymmetrically to replenish the major pool of less proliferative myoblastic Pax7-/ MyHC-transit amplifying cells (**TAC**) that are committed to differentiate ^21^.

To understand the potential impact of Ras isoforms on ciliation during the differentiation of C2C12 cells we applied mathematical modelling. We first estimated the fraction of asymmetric cell divisions in our system by generating an ordinary differential equation model (model 1) from our experimentally derived hierarchical model of C2C12 cell differentiation (**Figure S1A**). Using our previously published dataset of unperturbed C2C12 cell differentiation ^21^, model 1 predicted that 78 ± 1 % of MuSC divide asymmetrically. This fraction was in exact agreement with the experimentally determined fraction of ciliated Pax7+ cells (77.1 ± 7 %), which compares well to the 67 % ciliated MuSC isolated from murine muscle ^6^ (**Figure 1A,B**). By contrast, only 13 ± 2 % of the Pax7-transit amplifying cells (TAC) carry an Arl13B-positive primary cilium (**Figure 1A,B**), which again was similar to the 9.9 % reported in vivo ^6^. We further observed that in dividing ciliated C2C12 cells only one daughter cell re-assembles the cilium (**Figure S1B, Video S1**).

We therefore integrated ciliation as a factor that regulates asymmetric cell divisions of ciliated MuSC and TAC under differentiating conditions in a second model (model 2) (**Figure S1C**). Given that the majority of MuSC were ciliated (**Figure 1A,B**) and K-Ras depletion negatively impacted on the MuSC pool (**Figure S1D**) ^21^, we hypothesized that K-Ras promotes re-ciliation and thus protects stemness traits or restricts commitment during asymmetric cell divisions. We then applied model 2 to data describing how depletion of K-Ras4A/B, N-Ras or H-Ras perturb the differentiation of C2C12 cells (**Figure S1D**) ^21^. Fitting of the model 2 to these differentiation time-course data that report on the evolution of MuSC, TAC and differentiated cells (**Diff**) predicted that on day 3 of differentiation ∼27 % of cells remained ciliated (**Figure 1C**). By contrast, for data acquired after K-Ras4A/B ablation, a very low fraction of ciliated cells was predicted (∼1 %) (**Figure 1D**), while for N-Ras knockdown the fraction was similar (∼23 %) to the control (**Figure 1E**). H-Ras depletion on the other hand was predicted to increase the fraction of ciliated cells to ∼53 % (**Figure 1F**).

Experimental validation with previously validated siRNAs ^21^ and a K-Ras4B specific siRNA (**Figure S1E**), confirmed these predictions qualitatively for K-Ras4B (**hereafter K-Ras**) depletion, showing it reduced ciliation significantly, while the results of N-Ras depletion and H-Ras depletion were confirmed even quantitatively (**Figure 1G**). Loss of ciliation depletes MuSC during asymmetric divisions, while TAC increase thus promoting differentiation ^6^. In line with this, the above Ras-level manipulations led to an inverse impact on muscle cell differentiation that we assessed by myosin heavy chain (**MyHC**) expression (**Figure 1H**), confirming previous results ^21^. In further support of the K-Ras data, both expression of the dominant negative K-RasS17N (**Figure S1F,G**) or treatment with the pan-K-Ras inhibitor BI-2865, which also targets the wildtype protein, reduced ciliation while promoting differentiation (**Figure S1H,I**).

To summarize, these data suggest a Ras isoform-specific impact on ciliation and differentiation of C2C12 cells. Notably, K-Ras4B sustains re-ciliation and thus restricts differentiation by protecting in particular the MuSC pool from depletion during asymmetric cell division.

### Single-cell RNA sequencing of C2C12 cells identifies bona fide ciliated subpopulations

To embed our observations into the natural context of skeletal muscle cell differentiation, we performed single-cell RNA sequencing (scRNAseq). Samples were prepared to capture normal C2C12 cell proliferation and differentiation under high serum and low serum conditions, respectively, and the impact of the knockdown of K-Ras, transformation by K-RasG12C or knockdown of ciliary protein Ift88 on differentiation.

After quality control the resulting dataset consisted of 13,298 genes and 85,776 cells. We represented our sequencing results in a uniform manifold approximation and projection (UMAP) plot and performed clustering analysis (**Figure 2A**). Clusters were labelled as MuSC for the muscle stem cells in the cell line, TAC for committed myoblasts and myocytes and Diff for terminally differentiating myocytes and myofibers using established muscle differentiation genes ^22^ (**Figure 2B**).

**Figure 2.**
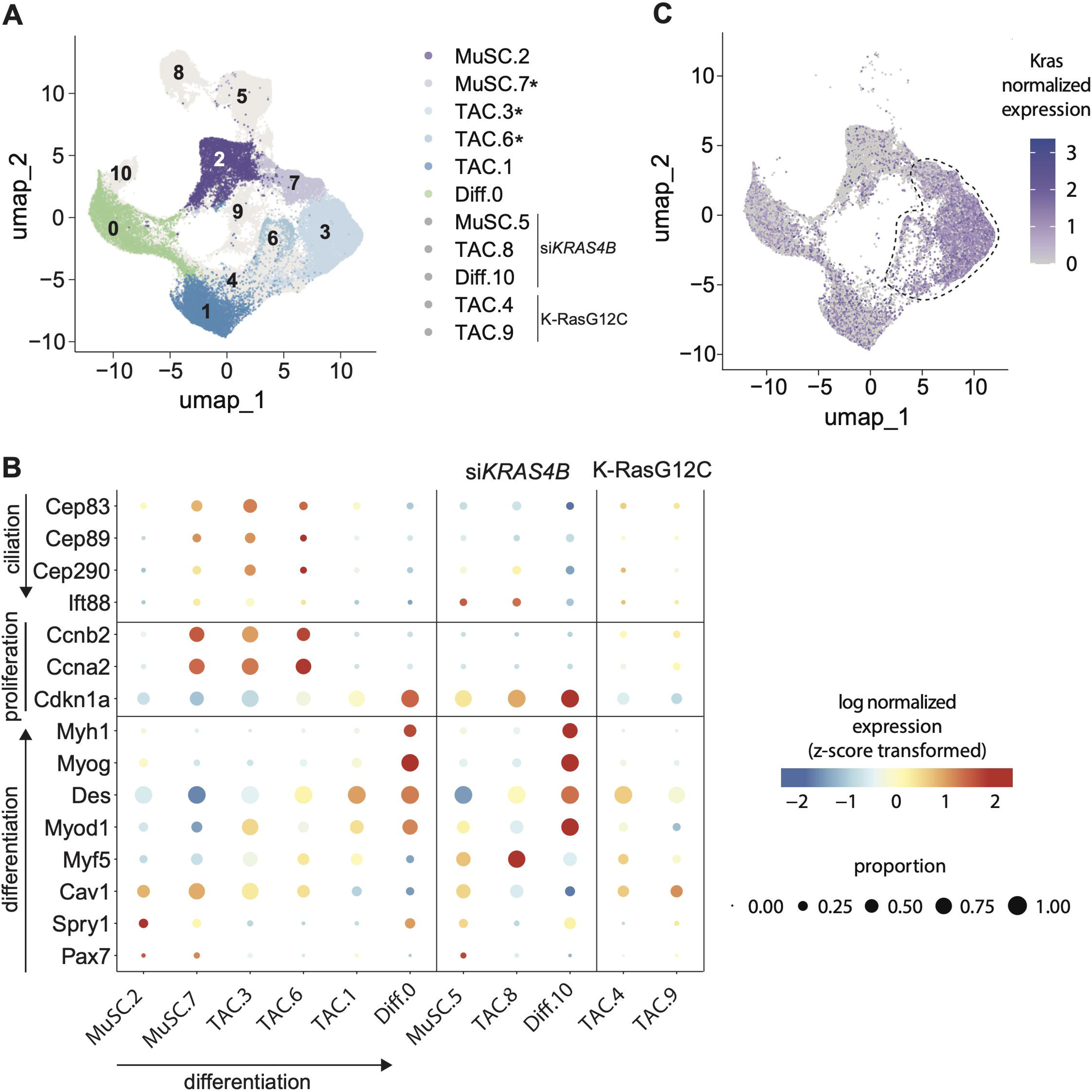
Modulation of K-Ras alters the C2C12 differentiation trajectory as identified by scRNAseq experiments. **(A)** UMAP representation of scRNAseq data with cell cluster labelling based on the expression of muscle differentiation marker genes as in (B). Shown in color are the cells that are associated with normal differentiation (high and low serum samples). Asterisks mark bona fide ciliated states. **(B)** Dot plots showing expression of marker genes in each cell cluster, as proportion of expressing cells (size) and gene-scaled expression level (color). Genes indicating ciliation and differentiation are ordered along the arrow toward maturation. Clusters with high expression of the satellite cell marker Pax7 were labelled as MuSC clusters. Pax7-negative clusters with elevated expression of Myf5 or commitment marker Myod1 comprised committed myoblasts and myocytes and were labelled as transit amplifying cell clusters (TAC). TAC represented the most heterogenous group of cell states in our samples. Clusters with high expression of differentiation markers Myog and Myh1 were labelled as Diff. Cluster numbering from the clustering analysis is preserved in the labelling. The ciliation gene set comprises Cep83 and Cep89, which regulate initial ciliary vesicle docking to initiate ciliary biogenesis ^63^. CEP290 participates during ciliary transition zone assembly in the next step, and IFT88 regulates ciliary elongation and protein transport within the fully assembled cilium ^64, 65^. **(C)** UMAP projection showing normalized expression of the *Kras* gene on a log scale in cells associated with normal differentiation (high and low serum samples). Bona fide ciliated cell clusters MuSC.7, TAC.3 and TAC.6 are marked with a dashed line.

We next inferred the putative unperturbed differentiation trajectory based on the well-established sequence of muscle differentiation marker expression, which was further supported by the Monocle 3 analysis (**Figure S2A-C**). This analysis was consistent with cluster MuSC.7 as the least differentiated, stem-like state, which progressed through TAC.3, TAC.6 and TAC.1 to the terminally differentiated cluster Diff.0 (**Figure 2B**). A potential alternative route via MuSC.2 directly to Diff.0 may be less significant given that MuSC.2 cells appear to be in a quiescent state (low Ccnb2, Ccna2 and high Spry1 expression) ^23^. This trajectory, including the alternative route to differentiation, aligns fully with a meta-analysis of in vivo derived data ^24^. Therefore, the C2C12 cell model is suitable to study cellular state transition processes during muscle cell differentiation.

To establish which clusters contain ciliated cells, we employed the predominantly ciliated Pax7+ MuSC subpopulation (MuSC.7) as a reference (**Figure 1A**). This subpopulation showed a high expression of genes relevant for early and late stages of ciliogenesis and an enrichment of cells expressing the late-stage structural protein of cilia Ift88 (**Figure 2B; Figure S2D**). In agreement with our imaging-based quantification of ciliation (**Figure 1B**), we inferred from the ciliation gene set that both Pax7-subpopulations TAC.3 and TAC.6 also contain ciliated cells.

Collectively, these data are in line with the existence of ciliated MuSC in vivo and the occurrence of several transitionally ciliated subpopulations as differentiation proceeds ^6, 22^.

### Modulation of K-Ras abundance or expression of oncogenic K-RasG12C perturb muscle cell differentiation

We noticed that all clusters MuSC.7, TAC.3 and TAC.6 with bona fide ciliated cell subpopulations had a relatively high number of *KRAS* expressing cells and were high in the expression of proliferation markers cyclin B2 and cyclin A2 (Ccnb2 and Ccna2) (**Figure 2B,C**). By contrast, *KRAS* and proliferation marker expression was lowest in non-ciliated cluster MuSC.2. The low serum condition associated cluster MuSC.2 may therefore represent a Pax7+/ Spry1+/ Cav1+ quiescent or reserve MuSC state, which under high serum conditions could reenter normal differentiation ^23^.

With the knockdown of K-Ras4B, clusters MuSC.5, TAC.8 and Diff.10 emerged, which expressed no proliferation markers but cell cycle inhibitor p21^Cip^^1^ (Cdkn1a) (**Figure 2B**). The former two clusters had only a low expression of genes encoding proteins required early in ciliogenesis, while still showing elevated Ift88 expression (**Figure 2B**). This may indicate that K-Ras4B ablation affects the abundance of cells that should become re-ciliated after cell division, consistent with our experimental observations (**Figure 1G**). Similarly, naturally low K-Ras4B expression appears to withdraw proliferative, ciliated MuSC.7 cells into the non-ciliated, quiescent MuSC.2 state. Artificial depletion of K-Ras4B induces then the MuSC.2-related states MuSC.5 and TAC.8 with an aberrant gene expression signature characterized by high expression of Pax7 and Myf5, respectively (**Figure 2B**).

Depletion of K-Ras4B furthermore led to the emergence of a new cluster Diff.10, which contained higher expression levels of terminal differentiation markers (**Figure 2B**). This can be explained by the fact that for terminal differentiation of muscle cells to occur the SPRED1/ NF1 tumor suppressor complex is engaged, which inactivates Ras ^25, 26^. Therefore, loss of K-Ras4B would also advance differentiation by this critical mechanism that appears to license terminal differentiation ^21^.

This is supported by our observations after transformation with oncogenic K-RasG12C. It is widely believed that oncogenic Ras mutations simply augment normal Ras activity and thus increase cell proliferation for instance to drive tumorigenesis ^27^. As such, one would expect the inverse of the cell state changes that are observed after K-Ras4B-knockdown, namely an advancement of cell differentiation (**Figure 2A**). However, the opposite is true, as oncogenic K-RasG12C appears to divert cells from the normal differentiation trajectory to halt them in the immature TAC.4 and TAC.9 states. These clusters have only low levels of proliferation markers in contrast to the high levels observed not only in clusters MuSC.7, TAC.3, but also TAC.6 under low serum/ differentiating conditions (**Figure 2B**). These findings confirm our previous observation that oncogenic Ras mutants block terminal differentiation and arrest cells in TAC-derived states with low proliferative activity ^21^, consistent with what is observed in mutant Ras-driven muscle borne cancer rhabdomyosarcoma (RMS) ^28^.

Altogether, scRNAseq data suggest that both K-Ras4B-depletion or expression of oncogenic K-Ras4B-G12C perturb normal differentiation but in different ways. Our data further support that a high K-Ras expression and activity may be required for re-ciliation in highly proliferative, ciliated subpopulations MuSC.7, TAC.3 and TAC.6. Thus, for differentiation to progress correctly, K-Ras activity may have to be up- or down-modulated depending on the cell state.

### K-Ras4B localization to the primary cilium of C2C12 cells depends on PDE6D

GTP-Arl3 localizes to the primary cilium where it functions as an allosteric release factor of high- and low-affinity clients of the trafficking chaperone PDE6D ^20^. We therefore expected that cancer-associated Ras proteins that bind with low, micromolar affinity to PDE6D are also released into the cilium ^14, 29^.

We transiently expressed GFP2-tagged wild type (wt) K-Ras (**Figure 3A**), N-Ras (**Figure 3B**) and H-Ras (**Figure 3C**) in C2C12 cells and examined their co-localization with ciliary marker Arl13B. K-Ras4A is not a cargo of PDE6D, due to constitutive steric clashes of its C-terminal residues at the entrance of the PDE6D prenyl-binding pocket and was therefore not examined ^14, 30^. Only K-Ras was significantly enriched in the cilium irrespective of its expression level, while N-Ras and H-Ras were less present (**Figure 3D**). This was expected, given that PDE6D engagement of N-Ras and H-Ras can be sterically blocked when they are reversibly palmitoylated near their prenylated C-terminal cysteine ^14, 15^. Consistent with a PDE6D-dependent delivery of K-Ras to the cilium, depletion of PDE6D essentially abrogated ciliary K-Ras localization (**Figure 3E,F**).

**Figure 3.**
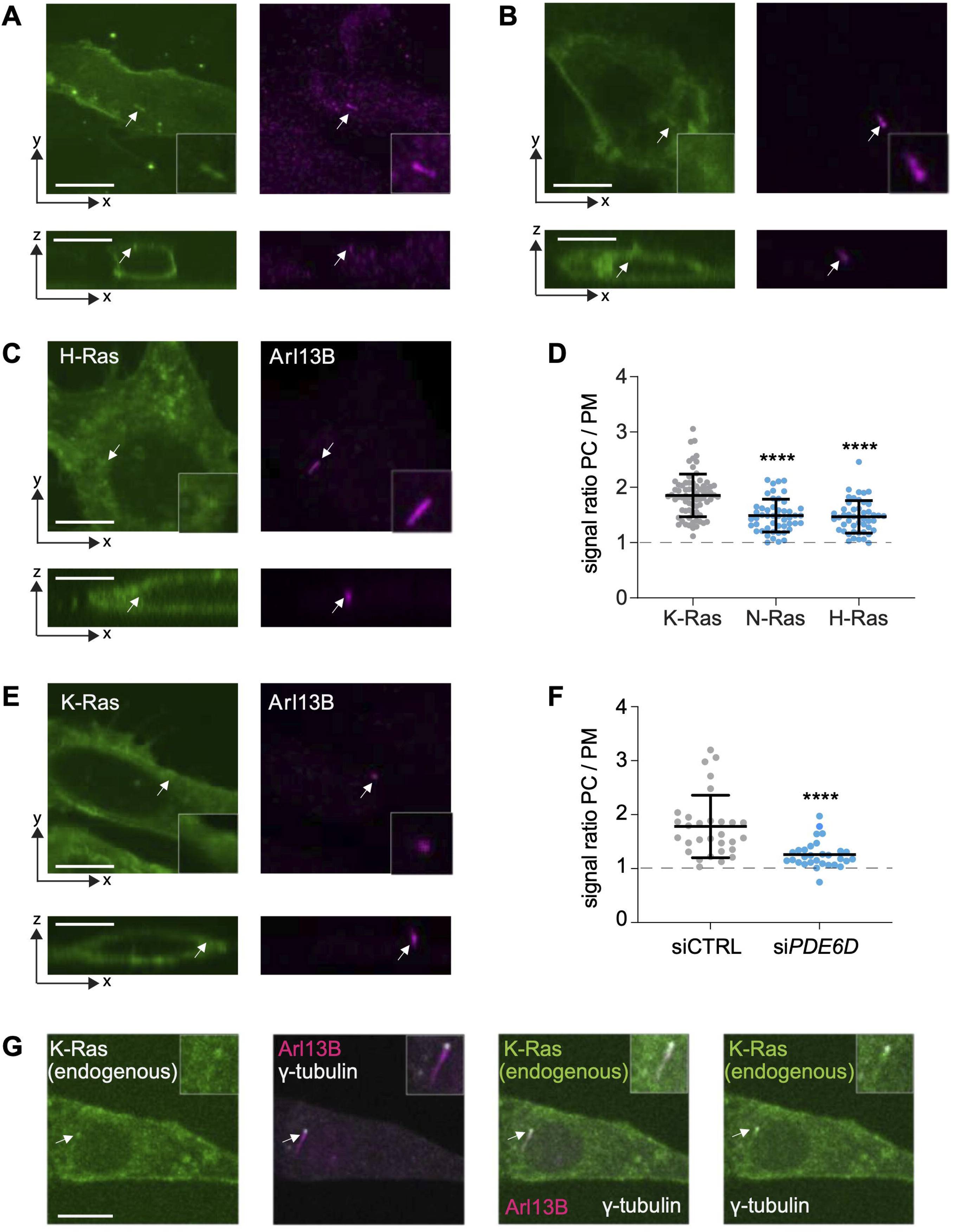
K-Ras accumulates PDE6D-dependently on the primary cilium. (**A-C**) Confocal images of C2C12 cells grown in high serum for 48 h after transfection with GFP2-K-Ras (A, n = 73), GFP2-N-Ras (B, n = 48) or GFP2-H-Ras (C, n = 47) and immunolabeled for ciliary marker Arl13B (arrow). *Bottom*, images show orthogonal views. Scale bar = 10 µm. **(D)** Quantification of Ras signal in the primary cilium (PC) as compared to the plasma membrane (PM) using data as in (A-C), N = 3. Means ± SD are plotted. Statistical comparison was done against K-Ras with the Kruskal-Wallis test and Dunn’s post hoc test. **(E)** Confocal images of C2C12 cells treated with PDE6D-targeting siRNA (100 nM), transfected with GFP2-K-Ras and cultured thereafter for 48 h in high serum. Scale bar = 10 µm. **(F)** Quantification of K-Ras signal in the primary cilium (PC) as compared to the plasma membrane (PM) using data as in (E), N = 3; siCTRL, n = 30; si*PDE6D*, n = 30. Means ± SD are plotted. Statistical analysis was done with the Mann-Whitney test. **(G)** Confocal images of C2C12-mEos3.2-KRAS knock-in cells cultured in high serum for 24 h directly after FACS-sorting and immunolabelled for ciliary marker Arl13B and centriolar marker γ-tubulin. Scale bar = 10 µm.

Using CRISPR-Cas9 genome editing of C2C12 cells we tagged the *KRAS* gene with the fluorescent protein mEos3.2. This allowed us to detect a faint signal of endogenous ciliary K-Ras at the plasma membrane and in the primary cilium (**Figure 3G**, **Figure S3A**). Transiently expressed mEGFP-K-Ras was furthermore observed in the cilium of serum starved MEF, NIH3T3 and hTERT RPE-1 cells (**Figure S3B-D**).

We conclude that amongst the cancer-associated Ras isoforms predominantly K-Ras localizes PDE6D-dependently to the primary cilium of C2C12 cells.

### Active K-Ras, B-Raf and active MEK appear to be confined to the centrioles at the base of the cilium

PDE6D facilitates diffusive trafficking of its cargo throughout the cell ^15^. However, the cilium represents a diffusion privileged reaction space, where only proteins smaller than ∼70 kDa can enter by diffusion, while larger ones need active transport ^11^.

To verify if K-Ras can enter the cilium by diffusion, we conducted Fluorescence Recovery After Photobleaching (FRAP)-experiments which confirmed that mEGFP-K-Ras (27 kDa tag + 21 kDa Ras) can move freely into the cilium (**Figure 4A**). Both the diffusion coefficient and mobile fraction were higher at the base as compared to the tip of the cilium (**Figure 4B-D**). Overall, diffusion appeared more than 10-fold reduced as compared to that of H-RasG12V determined previously in the bulk plasma membrane (D ≈ 0.4 μm^2^/s) ^31^. This may suggest that the ciliary tip can effectively act as a diffusion trap or reservoir for K-Ras.

**Figure 4.**
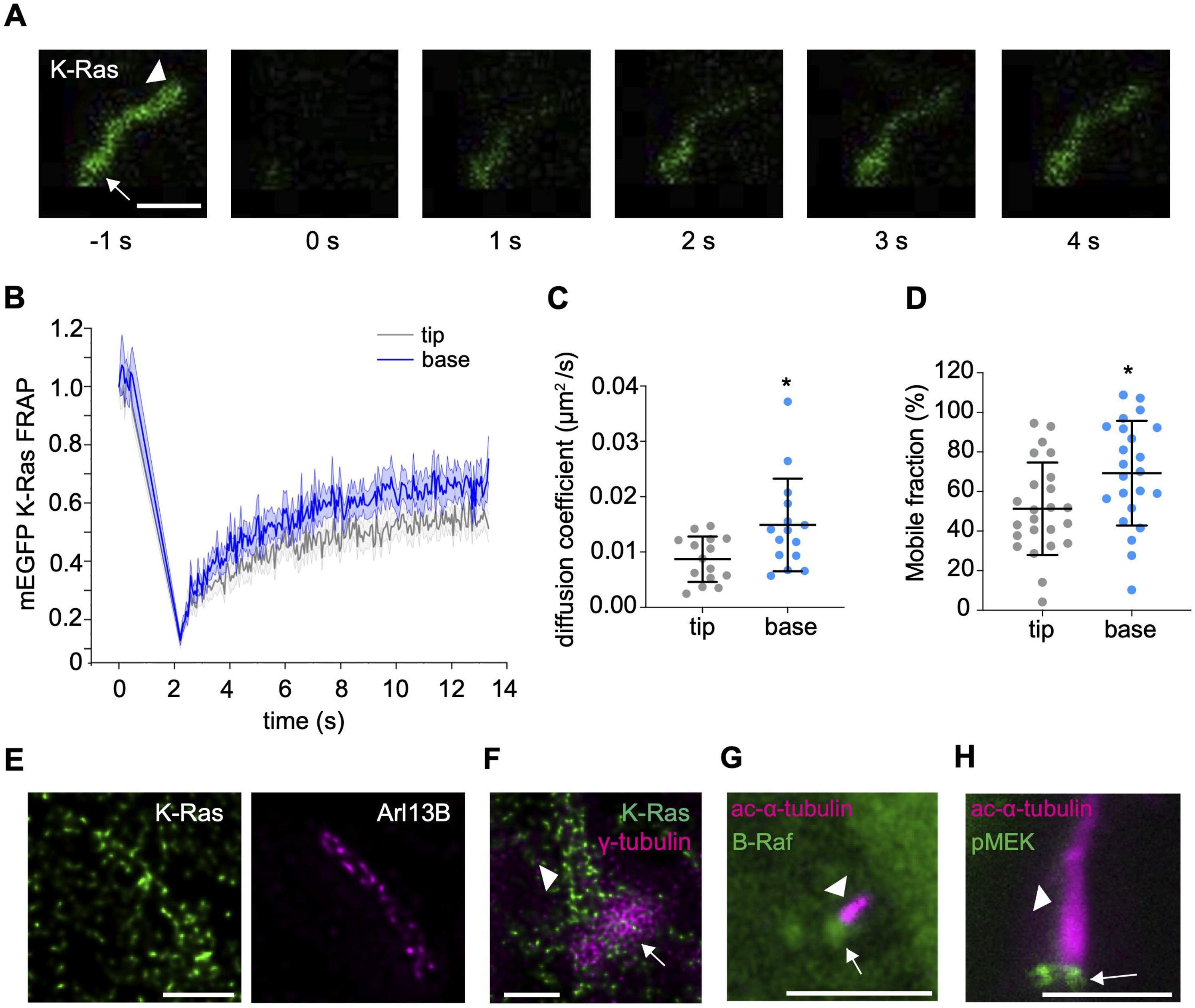
K-Ras localizes to the ciliary membrane while B-Raf and active MEK are confined to the centrioles. **(A)** Confocal images of C2C12 cells grown in high serum for 24 h after transfection with mEGFP-K-Ras. Images were acquired before (-1 s) and after (1 – 4 s) photobleaching at 0 s. Images represent the 10 µm × 10 µm photobleached region. FRAP was monitored at the ciliary tip (arrowhead) and wider base (arrow). Scale bar = 4 µm. **(B)** Recovery curves for tip and base using data as in (A). Mean ± SEM of fluorescence intensities from all cells per time point, N = 3, n = 23 per condition. (**C**, **D**) Analysis of individual FRAP curves summarized in (B) yielded diffusion coefficient (C; N = 3, n = 15) and mobile fraction (D; N = 3, n = 23) at ciliary tip and base. Means ± SD are plotted. Statistical analysis was done with the Mann-Whitney test. (**E, F**) STED images of C2C12 cells grown in high serum for 48 h after transfection with GFP2-K-Ras and immunolabeled for GFP using an ATTO647N-conjugated nanobody and ciliary membrane marker Arl13B or centriolar marker γ-tubulin. Position of the cilium (arrowhead) relative to the centrioles (arrow) is indicated. Scale bar = 2 µm. (**G, H**) Confocal images of C2C12 cells grown in high serum for 48 h. Cells were not transfected with K-Ras constructs, but with GFP2-B-Raf (G) and immunolabelled for acetylated α-tubulin marking the cilium (G; N = 3, n = 50) or immunolabelled for endogenous phospho-S217/S221-MEK1/2 and acetylated α-tubulin to mark the cilium (H; N = 3, n = 50). Scale bar = 2 µm.

Super-resolution STED-microscopy revealed that K-Ras distributed along the Arl13B-positive periphery of the cilium suggesting its association with the ciliary membrane (**Figure 4E**). In addition, GFP2-K-Ras appeared to be associated with the β-tubulin positive centrioles at the ciliary base (**Figure 4F, Figure S4A**). Consistent with its molecular weight, full length GFP2-tagged B-Raf (84 kDa B-Raf + 27 kDa tag) did not localize inside the cilium but was confined to the centrioles at the base of the cilium (**Figure 4G**). Likewise, the mCherry-tagged C-Raf-RBD (27 kDa tag + 11 kDa RBD), which binds only active Ras, was found only on the centrioles (**Figure S4B**), suggesting an accumulation of active Ras at that site. In line with MEK being phosphorylated in a complex with Raf ^32^, active S217/S221-phosphorylated endogenous MEK1/2 (45 kDa) was also only detected on the centrioles (**Figure 4H**). Active T202/Y204-phosphorylated endogenous ERK1/2 (42/44 kDa), however, appeared to localize throughout the cilium (**Figure S4C**).

In conclusion, K-Ras can freely diffuse into the cilium and localizes to the ciliary membrane and centrioles of the ciliary base. Essential MAPK pathway components (B-Raf, MEK1/2 and ERK1/2) localize and are active on the centrioles at the base of the cilium or inside the cilium.

### Ciliary K-Ras is sufficient to sustain normal ciliation and differentiation of muscle cells

To provide direct support that ciliary localization of K-Ras activity promotes ciliation and thus also restricts differentiation of C2C12 cells, we employed an artificial gain-of-function mutant. The K182S/K184I mutation immediately upstream of the farnesylated cysteine was inspired by the high-affinity, subnanomolar PDE6D cargo INPP5E and increases the affinity of K-Ras to PDE6D from micromolar to subnanomolar levels ^29^. K-Ras-K182S/K184I (**hereafter K-Ras-SI**) shows increased ciliary localization, while otherwise being sequestered to the nucleo-cytoplasm if sufficient PDE6D is available ^29^.

Consistently, the BRETtop value of K-Ras-SI with PDE6D was significantly increased as compared to the wild-type (**Figure 5A**) and effectively insensitive to S181-mutations in K-Ras-SI, which in the wild-type context significantly reduce the affinity to PDE6D ^14^ (**Figure S5A**). Furthermore, co-expression of PDE6D with K-RasG12C-SI shuts down the increase of pERK-levels observed otherwise in the absence of PDE6D in HEK cells, as expected by its quantitative sequestration to the nucleo-cytoplasm (**Figure S5B**) ^29^. By contrast, co-expression of PDE6D with oncogenic K-RasG12C significantly increased MAPK-output in HEK cells (**Figure S5B**), consistent with the notion that forward trafficking of K-RasG12C to the plasma membrane is enhanced by PDE6D and sustains bulk MAPK-signaling.

**Figure 5.**
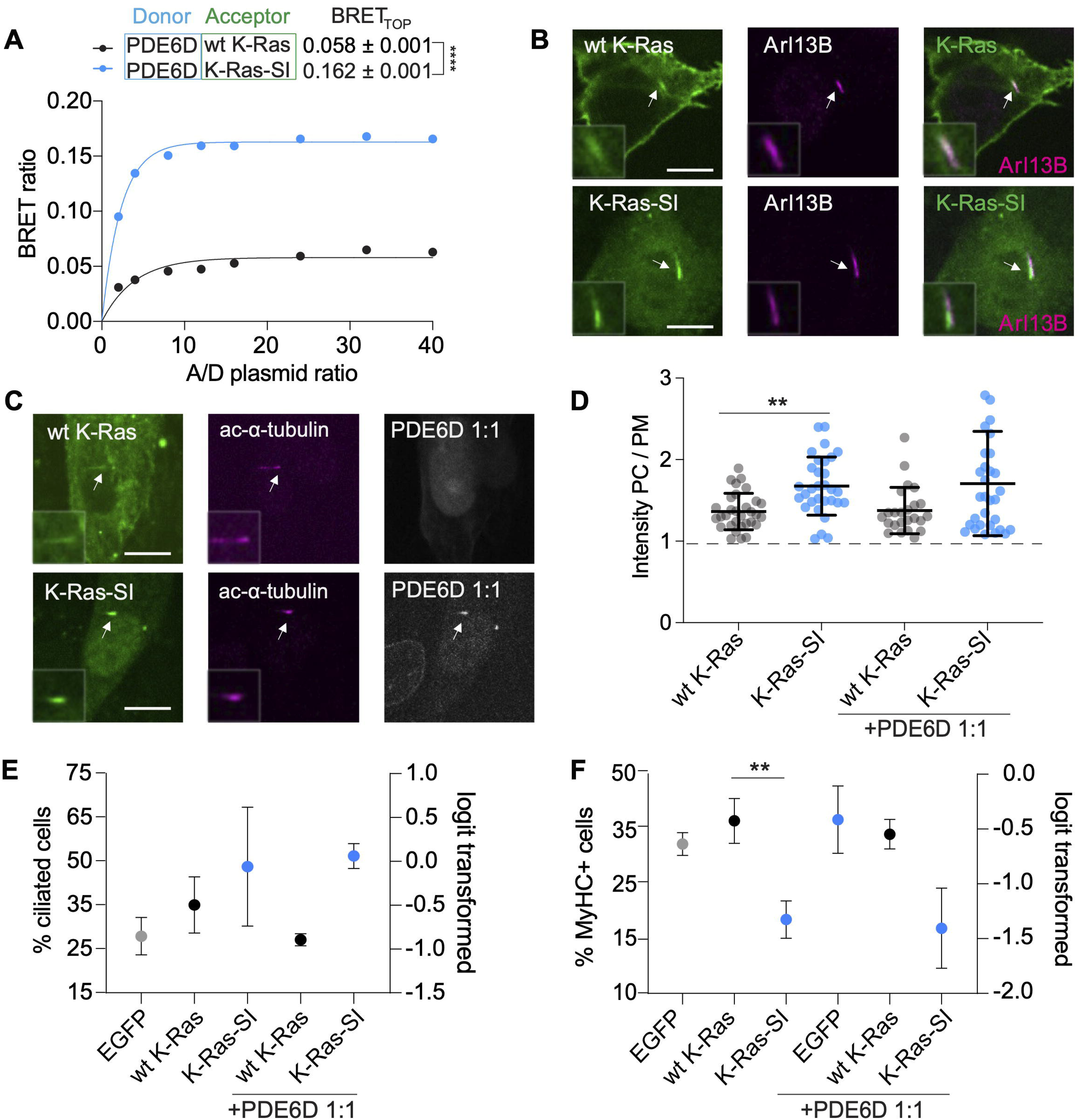
The ciliary localization gain-of-function K-Ras-SI mutant is sufficient to sustain ciliation and differentiation. **(A)** BRET titration curve of Rluc8-PDE6D and GFP2-K-Ras-SI as compared to GFP2-K-Ras expressed in HEK cells, N = 3. Means ± SD are plotted. Statistical analysis was performed with the extra sum-of-squares F-test. (**B, C**) Confocal images of C2C12 cells grown in low serum for 72 h after transfection with mEGFP-K-Ras (top) or mEGFP-K-Ras-SI (bottom) and immunolabelled for ciliary markers (arrows) Arl13B (B) or acetylated α-tubulin (C). The mCherry-PDE6D construct was co-expressed at 1:1 DNA ratio with K-Ras variant constructs (C). Scale = 10 µm. **(D)** Quantification of Ras signal in the primary cilium (PC) as compared to the plasma membrane (PM) per cell using data as in (B,C), N = 3, mEGFP-K-Ras, n = 31; mEGFP-K-Ras-SI, n = 31. Means ± SD are plotted. Statistical analysis was done with the Mann-Whitney test. **(E)** Confocal imaging-based quantification of ciliation of C2C12 transfected with the indicated mEGFP-tagged K-Ras constructs and cultured for 72 h in low serum. The mCherry-PDE6D construct was co-expressed at 1:1 DNA ratio with K-Ras variant constructs as indicated, N = 3, n ≥ 200 per condition. Means ± SD are plotted. Statistical analysis was done with one-way ANOVA and Dunnett’s T3 multiple comparisons test. **(F)** Flow cytometric quantification of MyHC terminal differentiation marker expression in C2C12 cells grown in low serum for 72 h after transfection with mEGFP-tagged K-Ras variant constructs alone or co-transfected with mCherry-PDE6D as in (E), N = 3. Means ± SD are plotted. Statistical analysis was done with one-way ANOVA and Dunnett’s T3 multiple comparisons test.

In our ciliated C2C12 cells K-Ras-SI localized to the cilium and nucleo-cytoplasm (**Figure 5B**). Co-expression of PDE6D enhanced this phenotype, while supporting quantitative nucleo-cytoplasmic sequestration to shut down any K-Ras signaling outside the cilium (**Figure 5C,D; Figure S5B**). In line with ciliary K-Ras promoting ciliation, K-Ras-SI expression increased ciliation and this effect was enhanced by co-expression of PDE6D (**Figure 5E**). Consistent with our Ras data (**Figure 1G,H**), differentiation with K-Ras-SI was significantly reduced as compared to wt K-Ras (**Figure 5F**). These data further support that ciliary K-Ras activity promotes ciliation and thus impacts on differentiation in our C2C12 cell system, as suggested by our mathematical modelling.

### K-Ras modulates ciliation-dependent heart-looping in zebrafish embryos

In order to appreciate, if K-Ras could more broadly regulate ciliation during development, we examined heart-looping in zebrafish embryos, which depends on the ciliated left-right organizer, called Kupffer’s vesicle in zebrafish ^33^. Thus, abnormalities in cilia morphology and/or function can lead to defects in heart-looping ^10^ (**Figure 6A**). Mechanical manipulation of cilia in Kupffer’s vesicle can facilitate normal or reversed heart-looping at will ^34^. Hence, heart-looping in zebrafish embryos is a read-out for cilia function in Kupffer’s vesicle *in vivo*. We therefore expressed wt K-Ras, K-Ras-SI and dominant negative K-RasS17N by micro-injection of synthetic mRNA in zebrafish embryos at the one-cell stage. Heart-looping was visualized at 48 hours post fertilization (hpf) by imaging the entire heart after *in situ* hybridization using a *myl7*-specific probe, which stains cardiomyocytes ^10^. Injection of neither wt K-Ras nor K-Ras-SI mRNA affected the proportion of embryos displaying normal and abnormal heart-looping (no looping and inversed looping) as compared to non-injected control embryos (**Figure 6B-D**). By contrast, K-RasS17N mRNA injected embryos showed reduced normal looping and a higher number of embryos with inverted looping and no looping (**Figure 6E**), consistent with significantly reduced ciliogenesis caused by this mutant in our muscle cell system (**Figure S1F**). These data illustrate that expression of dominant negative K-RasS17N affects left-right asymmetry, which is consistent with defective cilia formation and/or cilia function in Kupffer’s vesicle.

**Figure 6.**
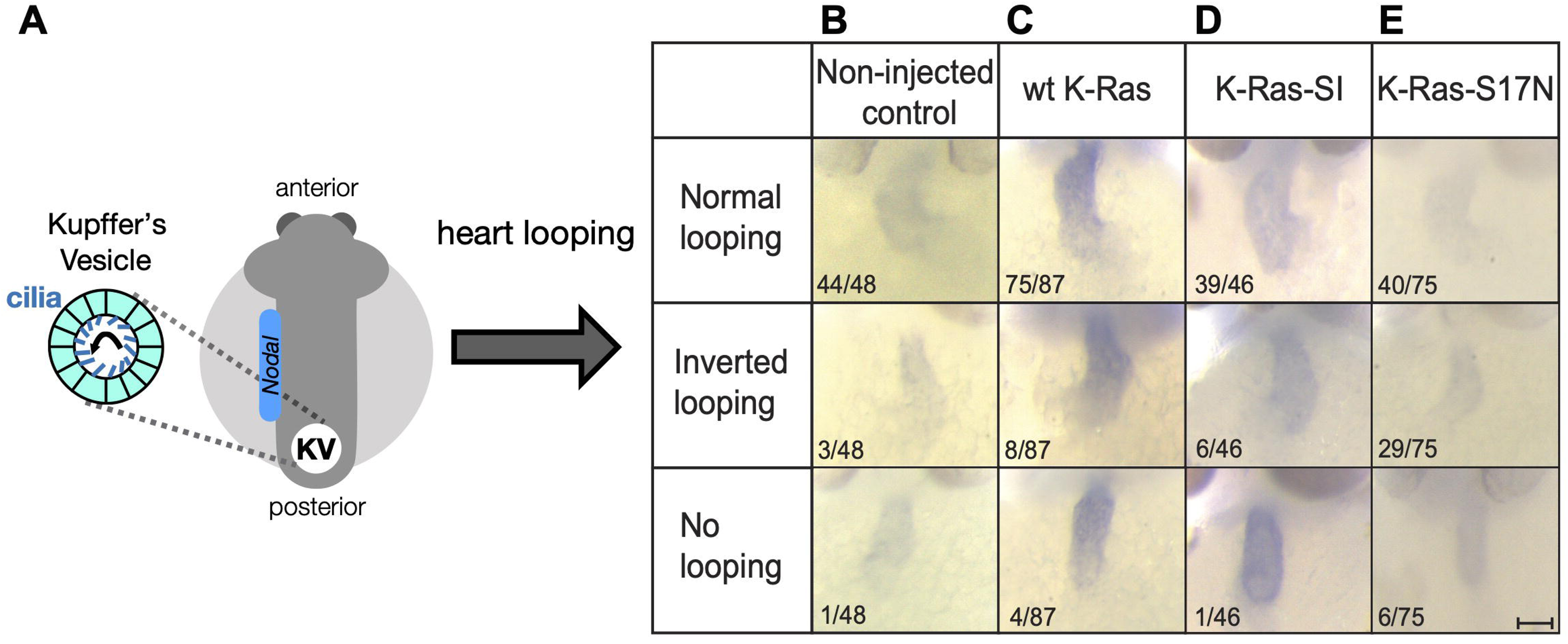
Modulation of ciliation-dependent heart-looping in zebrafish embryos by K-Ras mutants. **(A)** During gastrulation, body asymmetry is determined by a ciliated transient organ called Kupffer’s vesicle (KV) in zebrafish. Schematic shows dorsal view on embryo with region of asymmetrically expressed *Nodal* gene in blue. (**B-E**) Zebrafish embryos were not injected (Non-injected control, B), or microinjected at the one-cell stage with synthetic mRNA encoding wt K-Ras (C), K-Ras-SI (D) or K-RasS17N (E). The embryos were fixed at 48 hpf and *in situ* hybridization was performed with an RNA probe directed against cardiomyocyte-specific *myl7*. Images of representative embryos are shown here and the proportion of embryos displaying the specified heart-looping phenotype is indicated for each K-Ras variant. Scale bar = 100 µm.

In summary, modulation of K-Ras activity or abundance in the cilium may broadly impact ciliation-dependent differentiation and developmental processes.

## Discussion

We describe a novel fundamental role for K-Ras in the primary cilium of MuSC and TAC to restrict commitment or protect stemness during asymmetric cell divisions, which effectively reduces downstream differentiation. Access of Ras proteins to the primary cilium is mediated by the ciliary trafficking chaperone PDE6D, which prefers K-Ras over the other Ras isoforms for trafficking. This explains the predominant role of K-Ras in this context.

The exact sequence of molecular events and their spatio-temporal context downstream of ciliary K-Ras-MAPK-signaling that need to be enabled during asymmetric cell division remain at this point speculative. We propose that a high ciliary K-Ras-MAPK activity acts as or induces a ‘stemness mark’ that may become asymmetrically apportioned during cell divisions of ciliated cells. However, observing such processes directly is technically highly challenging. Endogenous K-Ras can be detected in cilia of C2C12 cells and after heterologous expression in three other mammalian cell lines, suggesting it can in general localize to this organelle. However, we describe and important functional role in C2C12 cells where it is involved in cell state changes as cells differentiate, while in the other ciliation model cell lines the formation of cilia is studied in isolation, typically after serum withdrawal. By contrast, in asymmetrically dividing C2C12-associated MuSC and TAC, ciliation and de-ciliation must occur in high serum as cells are integrated in a differentiation continuum. The developmental significance of K-Ras in ciliation processes is supported by our zebrafish experiments, which hint at its profound relevance during development.

How does K-Ras become activated in the primary cilium? Provided K-Ras can bind to PDE6D, it is plausible to assume that K-Ras is continuously delivered into the primary cilium, and its activation state there may thus reflect that of K-Ras in the whole cell. Additionally, K-Ras activity may be modulated by pathways resident in the cilium. Functionally, ciliary K-Ras may act as a reservoir, which sustains a high basal body activity of K-Ras. While it is possible that additional trafficking chaperones, such as CaM and centrin support trafficking of Ras to the cilium or basal body, their association probability with the here studied Ras isoforms follows similar rules as that with PDE6D ^35, 36, 37^.

Our mathematical model demonstrates that ciliation guides asymmetric cell divisions of C2C12 cells. It thus rationalizes how loss of ciliation after K-Ras depletion or inactivation can contribute to increased differentiation possibly by promoting the symmetrical division of a ciliated MuSC into two non-ciliated, committed TAC, instead of an asymmetric cell division. Interestingly, Ras isoform-specific effect on ciliation and differentiation were revealed, with N-Ras depletion having a neutral effect, while loss of H-Ras increased ciliation and decreased differentiation, while the opposite was seen for K-Ras. Additional effects of the Ras isoforms on differentiation downstream of ciliation can however not be excluded and require a dedicated study. We note that the qualitative behavior of the Ras isoforms observed here, strikingly correlates with that previously described in F9 mouse embryonal carcinoma stem cells ^38^. In those cells oncogenic K-RasG12V promoted proliferation of endodermal progenitors, while H-RasG12V was driving differentiation and N-RasG12V was neutral.

We observed loss of ciliation also with the expression of oncogenic K-RasG12C in TAC.9, consistent with the fact that most cancer cells do not carry a primary cilium ^39^. While not the focus of the current investigation, we want to discuss the ramifications of our observations for cancer. Our data suggest that the transforming activity of oncogenic Ras does not originate from increased proliferation, but from the differentiation block that is introduced in MuSC and/ or TAC, the stem and progenitor populations in the heterogenous C2C12 cell line. This finding contrasts with the fact that all major cancer cell assays and screens rely on cell proliferation or interpret Ras-MAPK-pathway activity in the context of proliferation ^40, 41, 42^. Instead, our data suggest shifting our attention to the specific Ras biology on centriolar organelles during differentiating cell divisions.

The C2C12 cells recapitulate the normal muscle cell differentiation program with relevant stem and progenitor cell subpopulations. This is supported by our scRNAseq data, which agree with a hierarchical continuum of myogenic stem- and progenitor states as observed in vivo ^43^. Transformed cells thus likely inherit the lineage properties, including the proliferation rate, which may explain why the natural frequency of stem cell divisions governs the susceptibility of tissues to develop cancer ^44^. Furthermore, transformed cells would compete against the background of tissue-resident stem cells, which as compared to terminally differentiated tissue then appears as hyperproliferation. These characteristics match exactly to those observed in Ras-driven RMS and more generally with the dedifferentiation and hyperproliferation hallmarks in cancer ^28^.

Given the here presented critical function of K-Ras to maintain stem- and progenitor cells, we predict that mutations of K-Ras in cancer are also deleterious, because they withdraw one or even two K-Ras alleles from the regulation of asymmetric divisions during normal tissue repair. This also explains the tumor suppressive activity of wildtype Ras as a function of keeping tissue healthy and normal stem and progenitor cells capable to not be outcompeted by their transformed counterparts ^45^. It is tempting to speculate that such a competition mechanism underlies the exploitation of the different Ras isoforms in cancer.

Due to the significant role of PDE6D in the cycle for the forward trafficking of K-Ras to the plasma membrane, it was previously explored as a surrogate drug target for K-Ras ^18^. However, given the importance of PDE6D in trafficking K-Ras to the cilium and thus to sustain stemness during asymmetric divisions, it should be regarded with caution as drug target. Based on our model, PDE6D inhibition would likely compromise the quiescence of stem cells, thus impacting on the regenerative potential residing in tissues. On the other hand, our data support that lowering its abundance could promote differentiation, which would in general be beneficial in the cancer context.

In analogy to the overactivation of Ras in cancer, it is currently believed that in RASopathies a milder hyperactivation of the Ras-MAPK pathway is the root cause ^46^. In this context the predisposition to cancer as observed in some RASopathies, goes back to the already elevated Ras-pathway activity. However, we propose that by controlling (Ras-MAPK-dependent) ciliation and thus cell differentiation, aberrant Ras-signaling perturbs tissue development and repair. When occurring locally in the adult, cancer can be promoted. If all cells of the body are affected, such as in RASopathy patients, tissue development is broadly affected ^1^. Despite the milder pathway activation as compared to those observed in cancer, differentiation defects become potentiated with each cell division round during development. Indeed, to be compatible with life, these pathway activations cannot be too extreme, in line with the fact that mutations occurring in cancer are typically not observed in RASopathies ^46^. By impacting on the formation of the cilium, which is the major developmental signaling hub, Ras furthermore affects multiple developmental and stemness signaling pathways ^47^. It is therefore plausible to assume that all these effects combined contribute significantly to the phenotypes observed in RASopathies.

Importantly, our model thus provides a molecular mechanistic explanation for the similar phenotypes observed in two groups of developmental diseases, RASopathies and multisystemic ciliopathies. It rationalizes how certain phenotypes of these two rare disease groups, such as intellectual disabilities, cardiovascular and skeletal abnormalities do not just overlap broadly, but originate in defects of the here described K-Ras-ciliation pathway. In line with this, others have already previously recognized the ciliary defects in RASopathy animal models ^9, 10^. Given the lack of in vitro models to study ciliopathies, we propose that the C2C12 cell line could also here serve as a simple model system for this disease group. Moreover, RASopathy treatments benefit significantly from developments in oncology, and with the newly found commonalities with ciliopathies, Ras-MAPK pathway inhibitors may find an application in the treatment of ciliopathies.

In conclusion, our study suggests a major role of Ras proteins to guide cell differentiation with a major role of K-Ras in the primary cilium. Given that most stem and progenitor cells are ciliated and the molecular machinery that actuates ciliary K-Ras is conserved in most cell types, it can be assumed that the above mechanism applies widely during organismal development. In addition to developmental diseases and cancer, we therefore foresee a broad impact of our mechanism on stem cell, aging and regenerative research.

## Materials and methods

### Materials and equipment

A list of employed materials and equipment with their sources and identifiers can be found in Table S1.

### Cell lines

All cell lines except hTERT RPE-1 were maintained in Dulbecco’s Modified Eagle’s Medium (DMEM) supplemented with ∼9 % (v/v) fetal bovine serum (FBS), 2 mM L-glutamine and 1 % penicillin/streptomycin. In the context of C2C12 cell culture, this medium is referred to as ‘high serum medium’. For culture of hTERT RPE-1 cells, DMEM/ F-12 supplemented with ∼9 % (v/v) fetal bovine serum (FBS), 2 mM L-glutamine, 1 % penicillin/streptomycin and 0.01 mg/mL hygromycin B (RPE-1 medium) was used. Cells were incubated in a humidified incubator at 37 °C, with 5 % CO_2_ and passaged twice per week at a confluency of 50-60 %. C2C12 differentiation assay was carried out as described previously ^21, 48^. Briefly, C2C12 cells were first allowed to proliferate up to ∼ 90 % confluency. The ‘high serum’ cell culture medium was then exchanged with ‘low serum medium’, which is DMEM supplemented with 2 % horse serum. Fresh low serum medium was added every day for three days. Specific treatment conditions are indicated in the figures.

### Expression constructs

Multi-site gateway cloning was utilized to prepare expression constructs of most Ras variants, except K-Ras S17N, and to prepare B-Raf, and PDE6D constructs. If gene variants were obtained as g-block fragments, BP clonase II enzyme mix was used to recombine g-blocks with pDONR221 donor plasmid to create entry clones with flanking sites compatible with LR cloning. Variant entry clones were recombined with entry clones containing the CMV promoter and entry clones containing Rluc8, GFP2, mEGFP or mCherry into the pDest305 or pDest312 destination vectors ^49^. LR clonase II enzyme mix was utilized to perform LR Gateway recombination and thereafter the recombination mix was transformed into ccdB-sensitive *E.coli* strain DH10B. Ampicillin resistant colonies were screened for the expressed plasmid via restriction digestion with BsrGI-HF. To generate the pEGFP-K-Ras S17N construct, mEGFP was cloned into the vector backbone pcDNA(3.1)-mCherry K-Ras S17N through restriction digestion with enzymes NheI and BsrGI-HF. K-Ras variants used for mRNA-transcription and injection into zebrafish embryos were cloned into pCS2+ vectors using a one-step isothermal Gibson assembly. All final constructs were validated with Sanger sequencing.

### Cell lipofection

C2C12, MEF and NIH-3T3 cells were transfected with plasmid constructs as indicated in the figure legends, using Lipofectamine 2000. Cells cultured in high serum were transfected at a confluency of 50-60 %. Plasmid DNA amounts were as per manufacturers’ guidelines. Briefly, 2 µg plasmid DNA was diluted in 125 µL Opti-MEM medium and mixed by vortexing for 2-3 s. This mixture was incubated at 23-25 °C for 5 min. Simultaneously, 7 µL Lipofectamine 2000 reagent was added to 125 µL Opti-MEM medium and mixed by vortexing for 2-3 s. This mixture was incubated at 23-25 °C for 5 min as well. Thereafter, Lipofectamine-Opti-MEM mixture was added to the plasmid DNA-Opti-MEM mixture, vortexed for 2-3 s and incubated for 10 min at 23-25 °C. This solution was then added dropwise to one well of a 6-well plate containing cells in 2 mL high serum medium. After 4 h the medium was replaced with fresh high serum medium. To induce differentiation, the medium was exchanged with low serum medium 24 h after transfection. Low serum medium was replaced every day for three days.

### Nucleofection of hTERT RPE-1 cells

Lonza 4D-Nucleofector X unit with the SE cell line nucleofection kit was used to deliver plasmid DNA into hTERT RPE-1 cells. Nucleofection was carried out as per the manufacturer’s instructions. Briefly, cell passaging was performed when the cells reached 80-90 % confluency and 1 mL aliquots of cells at a concentration of 1 × 10^6^ cells/ mL were prepared in RPE-1 medium. After centrifugation at 90 × *g* for 10 min at 37 °C, the supernatant was discarded and the cell pellet was resuspended in 100 µL nucleofection solution supplied with the SE cell line nucleofection kit. Plasmid DNA at a concentration of 2 µg was added to this cell suspension. The cell suspension was then transferred to the bottom of a nucleofection cuvette supplied with the SE cell line nucleofection kit. The cuvette was added to the retainer of the 4D-Nucleofector X unit and nucleofection was carried out with the DS150 program. The cuvette was thereafter removed from the retainer of the nucleofector unit and after pipetting gently, the cell suspension was added to a 6 well plate containing RPE-1 medium pre-warmed at 37 °C. Medium was switched to RPE-1 medium without FBS 24 h later and cells were incubated with the serum free medium for another 24 h to induce ciliation.

### siRNA transfection

For siRNA transfections in 6-well plates, typically 100 nM siRNA was diluted in 250 µL Opti-MEM transfection medium containing 7.5 µL Lipofectamine RNAiMAX reagent. Transfection was performed when cells grown in high serum culture medium reached 50-60 % confluency. The transfection mix in Opti-MEM was then added to 2 mL high serum medium per well of a 6 well plate. This medium was subsequently replaced 24 h later with low serum medium. Low serum medium was replaced every day for three days.

### Immunofluorescence

C2C12 cells were seeded on coverslips with a thickness of 0.17 mm in 6-well plates at 100,000 cells/ mL. Cells were transfected 24 h later with the indicated fluorescent protein tagged constructs and cultured for 48 h in high serum medium or 72 h in low serum, as stated in the figure legends. Cells were then fixed with 4 % (w/v) paraformaldehyde in PBS and permeabilized with 0.5 % Triton X-100 for 10 min. Then, cells were washed with 0.05 % Tween 20 in PBS (PBST) and incubated 30 min with 2 % BSA in PBST. Specific antigen labelling was done for 1 h using primary antibodies. Cells were subsequently washed 3 × 5 min with PBST, followed by 1 h incubation with secondary antibodies or with Phalloidin-Alexa Fluor 647. After washing 5 min with PBST, nuclei were counterstained with 0.2 μg/ mL Hoechst 33342 in PBST and washed again with PBST. A drop of Vectashield mounting medium was added on glass slides and the coverslips were mounted on them.

### Confocal imaging and quantification of ciliation

Fixed cells on coverslips were imaged using a 60 × NA 1.3 oil immersion objective on a Nikon Ti-E microscope equipped with a Yokogawa CSU-W1 spinning disk confocal unit and an Andor iXon Ultra EMCCD camera. GFP2-, EGFP- or mEGFP-tagged constructs were visualized by excitation with a 488 nm laser line and detected using EGFP emission settings with a 535/20 band pass filter. Excitation of mCherry or Alexa Fluor 594 was with the 561 nm laser line and Alex Fluor 647 excitation was achieved with the 640 nm laser line. Alexa Fluor 594 and mCherry were visualized with the 560/40 band pass filter and Alexa Fluor 647 was visualized with the 700/35 band pass filter. Z stacks were acquired with 0.3 μm spacing between each optical section. Images were acquired with Nikon NIS-elements software and analyzed in Fiji/ ImageJ ^50^.

To quantify relative abundance of Ras in the cilium, 3D orthogonal views were obtained for each image using Fiji. The intensity of GFP2- or mEGFP-tagged Ras-constructs was measured on the primary cilium (PC) in the orthogonal stacks, as the average signal of a region of interest around the cilium identified by labelling the ciliary marker Arl13B. The intensity of Ras at the plasma membrane (PM) was the average signal of a region of interest around the entire PM. Subsequently the relative ciliary localization was calculated by dividing the two averaged signals as PC/PM-signal ratio.

For quantification of ciliated cell fractions, images were acquired to visualize Hoechst 33342-labelled nuclei and cilia marked with anti-Arl13B antibody detected with Alexa Fluor 647-conjugated secondary antibody. For each image, nuclei and cilia were manually counted. The percentage of ciliated cells per image was thus obtained by calculating the proportion of ciliated cells relative to the total number of nuclei. At least three images containing at least 100 cells were quantified in this manner per biological repeat and the average of percentages thus obtained was plotted as the ‘% ciliated cells’.

For live cell imaging, C2C12 cells were seeded at a density of 100,000 cells/mL in high serum medium in glass bottom petri-dishes and transfected 24 h later with pEGFP-C1-Centrin and pDest312-CMV-Arl13B-mCherry plasmids. Medium was replaced with fresh high serum medium 4 h after transfection. After a further 48 h, the medium was replaced with imaging medium (DMEM high glucose, without glutamine and phenol red, supplemented with ∼9 % FBS). Cells were imaged with an Andor BC3 benchtop confocal microscope equipped with a plan fluor 40 × NA 0.95 objective, at 37 °C with 5 % CO_2_. EGFP-Centrin was visualized by excitation with a 488 nm laser line and detected using EGFP emission settings with a 535/20 band pass filter. Excitation of Arl13B-mCherry was performed with 561 nm laser line and it was visualized with the 560/40 band pass filter. Time-lapse Z-stacks, with 0.3 μm spacing between each optical section were acquired at intervals of 5 min for a total period of 24 h. Images were analyzed with Fiji and presented as collapsed Z-stacks representing each time point.

### STED microscopy

To visualize K-Ras with STED microscopy, C2C12 cells expressing GFP2-K-Ras were fixed with 4 % (w/v) paraformaldehyde and immunolabelled with ChromoTek GFP-booster ATTO647N. To visualize primary cilia, cells were immunolabelled with primary antibody against Arl13B (177-11-1AP, Proteintech; dilution 1:500) and goat anti-Rb Alexa Fluor 594 secondary antibody (cat. no. A-11012, ThermoFisher Scientific; dilution 1:200). Samples were mounted on glass slides with Vectashield mounting medium.

The STED imaging was performed using the Abberior Infinity Line microscope (Abberior GmbH, Göttingen, Germany) equipped with Imspector software version 16.3.16118-w2224 and an Olympus 60 × oil objective (1.42 NA). To obtain two-color STED images, the fixed cell samples labelled with ATTO 647N and Alexa Fluor 594 dyes were excited with 640 nm and 561 nm picosecond pulsed laser diodes at 40 MHz (Abberior GmbH, Göttingen, Germany), respectively. STED imaging was conducted using the 775 nm pulsed STED laser (NKT Photonics Switzerland GmbH, Regensdorf, Switzerland) at 40 MHz to capture the desired STED images for both fluorophores. Emitted light was detected by using an avalanche photodiode detector (Excelitas Technologies Corp., USA) with 5 µs dwell time and line averaging of 10. To avoid cross-talk between the two channels, the samples were initially excited exclusively with the 640 nm laser at 16.7 µW power, while the 775 nm STED laser at 177.6 mW power was used to acquire STED images of ATTO 647N-labelled samples. After a brief interval of 1-2 min, the second channel was activated, employing the 561 nm laser at 3.42 µW power and the 775 nm STED laser at 365.9 mW power, to obtain STED images of Alexa Fluor 594-labelled samples. In this study, the detection range for ATTO 647N was set from 650 nm to 754 nm, while for Alexa Fluor 594, the detection range was set from 584 nm to 713 nm. These ranges were chosen to ensure optimal detection of fluorescence signals emitted by the respective dyes. Images were exported at 16-bit in the Abberior-MSR format.

Following data acquisition, Huygens professional software version 23.4 was employed for STED image deconvolution. Our experimental imaging parameters were filled into the software, and subsequently, the Huygens Deconvolution Wizard was utilized, offering the option to use either the measured or theoretical point spread function (PSF). The theoretical PSF, based on our experimental imaging parameters, was generated within the Huygens software. Additionally, the software automatically estimated the background and signal-to-noise ratio for each image, facilitating their utilization in the Classic Maximum Likelihood Estimation algorithm for generating deconvolved images.

### Fluorescence recovery after photobleaching (FRAP)

C2C12 cells were seeded at a density of 100,000 cells/ mL in high serum medium in a coverslip-bottom dish and co-transfected 24 h later with pDest-305-CMV-mEGFP-K-Ras4B and pDest305-CMV-Arl13B-mCherry plasmids. Medium was replenished with fresh high serum medium after 4 h. After a further 24 h incubation, medium was exchanged with imaging medium (DMEM high glucose, without glutamine and phenol red, supplemented with ∼9 % FBS) and cells were then observed on a Zeiss LSM 980 confocal laser scanning microscope equipped with a Plan-apochromat 40 × water immersion objective (NA 1.2) and GaAsP hyperspectral detectors. The cilium was identified by the presence of Arl13B-mCherry whose fluorescence was detected with excitation using the 561 nm laser line and detection around 610 nm i.e. standard mCherry detection settings in the Zeiss LSM software. Ciliary localization of mEGFP-K-Ras was verified by excitation with the 488 nm laser line and detection via standard EGFP settings. Pre-bleach images of EGFP-K-Ras were acquired at low laser power (6 μW) of the 488 nm line. Photobleaching of mEGFP-K-Ras was performed by setting the 488 nm laser line at the maximum possible laser power (2 mW) with 110 bleaching iterations of 0.5 ms in a 10 μm × 10 μm rectangular region encompassing the cilium. Fluorescence recovery at the broad cilium base and the narrow, tapering cilium tip was monitored by setting up circular ROIs of radius 0.4 μm. Serial micrographs of resolution 512 × 512 pixels were acquired at the maximum scan speed without averaging for a period of 13.5 s at intervals of 0.5 ms. Fluorescence intensities from both ROIs were exported in CSV files from Zeiss Zen Blue software and post-bleaching intensities were plotted as a fraction of pre-bleach fluorescence intensity. Fluorescence recovery curves were generated in OriginPro, which represent the mean fluorescence intensities ± SEM obtained from all cells at each time point.

For calculating the diffusion coefficient of fluorescent molecules in the circular ROI regions, the following equation was used ^51^:

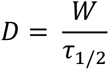

*W* represents the ROI radius and *τ_1/2_* denotes the half-time, indicating the time point when the fluorescence intensity equals half of the fluorescence intensity at maximum recovery, here the experiment end-point. Calculation of *τ_1/2_* was done by fitting the fluorescence recovery curve to a first-order fitting function using a python algorithm ‘Fitting-curves-python’. The fitting function is as follows:

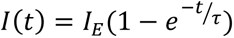

Where *I(t)* is the fluorescence intensity at time *t*, *I_E_* is the fluorescence intensity after full recovery and *τ* is the characteristic recovery time constant. *τ_1/2_* can then be calculated as follows:

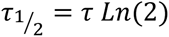

To calculate the immobile and mobile fractions of fluorescent molecules in the ROI, the following equation was used:

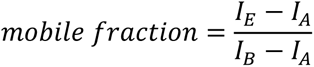

Where *I_B_* is the fluorescence intensity before the photobleaching step, *I_A_* is the fluorescence intensity immediately after the photobleaching and *I_E_* is fluorescence intensity after full recovery i.e. at the end-point of the measurement. Mobile fraction values thus obtained were multiplied by 100 and expressed as percentage of mobile fraction at the tip and base.

### Flow cytometry-based C2C12 cell differentiation assay

The differentiation assay was performed on a flow cytometer Guava easyCyte HT-2L flow cytometer as described by us previously ^21, 48^. Typically, > 1000 cells were analyzed in the final target gate per experimental repeat. In brief, C2C12 cells were seeded at a density of 100,000 cells/ mL in 6 well plates and treated as indicated in the figure legends. Low serum medium with drugs was replaced every day for three days. Cells were then harvested by trypsinization for 5 min and pelleted by centrifugation at 500 × g for 5 min. The cell pellet was fixed with 4 % (w/v) formaldehyde in PBS for 10 min. After washing with PBS, cells were permeabilized with 0.5 % Triton X-100 in PBS for 10 min. Subsequently cells were washed with PBST and immunolabelled with eFluor 660-conjugated anti-myosin 4 MF20 (myosin heavy chain/ MyHC antibody detecting the Myh1 gene product), diluted 1:100 in PBST for 1 h at 4 °C. Cells were pelleted by centrifugation at 500 × g for 5 min and resuspended in PBST for flow cytometric analysis. Gates and detection channel settings were established using non-labelled, EGFP-only expressing and MyHC-positive cells, as described ^48^. Intact cells were analyzed for the expression of EGFP in the GFP-low bin (up to 10-fold above background fluorescence) and MyHC. EGFP-variants were detected by 488 nm excitation and the Grn-B (525/30) band pass filter. MyHC immunolabelled with eFluor 600-conjugated antibody was detected by 640 nm excitation and the Red-R (664/20) band pass filter. To analyze MyHC-positive cell fractions in cells expressing mCherry-PDE6D, data was acquired on a BD LSRFortessa flow cytometer which was equipped with 405 nm, 488 nm, 561 nm and 640 nm laser lines. EGFP-variant fluorescence was detected by 488 nm excitation and the 530/30 bandpass filter. MyHC positive cells labelled with eFluor 660 and mCherry positive cells were excited with the 640 nm and 561 nm laser lines, respectively, and emission was recorded using the 670/14 and 610/20 band pass filters, respectively. Differentiation was quantified as the percentage of MyHC-positive cells in the GFP-low bin. Quantification was performed using FlowFate. For samples with mCherry-PDE6D, only mCherry-positive cells were considered for analysis and MyHC-positive cells in the GFP-low window were quantified manually with FlowJo.

### Fluorescence-activated cell sorting (FACS)

C2C12-mEos3.2-KRAS knock-in cells were provided as a heterogenous pool by Synthego Inc. where the CRISPR-Cas9 edited mEos3.2 KRAS knock-in cells only comprised a minor fraction of the total cell pool. Cells harboring the KRAS knock-in were enriched by cell sorting with a BD FACS Melody instrument equipped with 405 nm, 488 nm and 561 nm laser lines. Cells were grown in 175 cm^2^ flasks and upon reaching ∼90 % confluency, were detached by trypsinization, centrifugated at 1000 × *g* for 3 min and resuspended in 5 mL FACS buffer (PBS + 1 % FBS). mEos3.2-positive cells were identified through excitation with the 488 nm laser line and detection with the 530/20 band pass filter. The baseline was established with 488 nm excitation of non-fluorescent parental C2C12 cells lacking any mEos3.2 expression. Cells were sorted into 15 ml tubes containing 1 mL high serum culture medium, and at least 100,000 cells were collected at each sorting session. Sorted cells were subjected to centrifugation at 1000 × *g* for 3 min, resuspended in 2 mL high serum culture medium and plated into one well of a 6 well plate. After 24 h culture, cells were fixed with 4 % (w/v) PFA and processed for immunofluorescence labelling.

### Zebrafish heart-looping analysis

For the expression of K-Ras variants in zebrafish embryos, mRNA was synthesized using the mMessage mMachine SP6 transcription kit using the plasmids pCS2+-CMV-mEGFP-K-Ras4B-wt, pCS2+-CMV-mEGFP-K-Ras4B-SI and pCS2+-CMV-mEGFP-K-Ras4B-S17N as templates. Tüpfel longfin zebrafish embryos were collected for microinjection at the one-cell stage ^52^. The amount of mRNA that was injected per embryo was titrated down to the amount of mGFP-K-Ras-wt that did not induce morphological defects. This amount of mRNA (65 pg per embryo) was injected for all K-Ras variants. The injection mixes containing mRNA were run on a 1 % TAE agarose gel to verify the mRNA concentrations of the three K-Ras variants. Embryos were maintained in E3-medium. To inhibit pigment formation, embryos were treated with 1-phenyl 2-thiourea at 24 hpf. At 48 hpf, embryos were dechorionated using tweezers and fixed in 4% PFA in PBS at 4 °C overnight followed by a gradual dehydration to methanol. Embryos were stored in methanol at -20 °C. *In situ* hybridization was performed using a cardiomyocyte-specific RNA probe directed against *myl7* ^53^. Embryos were mounted in 2% methylcellulose and embryos were imaged using a Leica M165FC microscope with a Leica DMC5400 camera.

### BRET assay

BRET assays were performed following our previously described protocol ^37, 54^. Briefly, 200,000 cells/ mL HEK293 EBNA cells were plated and maintained for at least 72 h in 12-well cell culture plates. After 24 h, a donor construct tagged with Rluc8 and an acceptor construct tagged with GFP2 were transfected into cells using 3 μL of jetPRIME transfection reagent. BRET-titration experiments for donor saturation were performed by transfecting a constant concentration of donor plasmid (25 ng) and an increasing concentration of acceptor plasmid (from 0 to 1,000 ng). An identical total DNA load of 1,050 ng per well was achieved by adding pcDNA3.1 plasmid without insert. Cells were cultured for 48 h and prepared for the BRET assay. A CLARIOstar plate reader was used to perform BRET-measurements at 25 °C as described ^55^. Four technical replicates were measured for each experimental condition using signal channels that are specific for the luminophores. The GFP2-acceptor signal was recorded at λex = 405 ± 10 nm and at λem = 515 ± 10 nm. After addition of 10 μM coelenterazine 400a simultaneous recordings of Rluc8-signals for donor signal λem = 410 ± 40 nm and for the BRET-signal at λ = 515 ± 15 nm was performed. The calculation of the BRET ratio was done as previously described ^55^.

For BRET donor saturation titration experiments, the BRET ratio was plotted against the acceptor/donor plasmid ratios. All data typically averaged from three biological repeats were plotted at once using nonlinear regression models and were fitted into one phase association equation of Prism 9 or 10 (GraphPad). The Y_MAX_ value, which represents the top asymptote on the Y-axis, was taken as characteristic BRET_TOP_ value, which represents the maximal BRET ratio reached within the defined acceptor/donor ratio range. If no plateau can be recognized in the data, we did not use the fitting but instead state the value of the highest BRET ratio reached at the highest acceptor/donor ratio as BRET_TOP_.

### Immunoblotting

Prior to transient transfection, 250,000 HEK293 c18 cells were seeded in 1 mL DMEM per well of 12-well plates. After 24 h, cells were transfected with 1 µg of plasmid DNA and 3 µL Lipofectamine 2000 in Opti-MEM medium.

In situ cell lysis was performed 24 h after transfection as described before in ice-cold lysis buffer (50 mM Tris-HCl pH 7.5, 150 mM NaCl, 0.1 % w/v SDS, 5 mM EDTA, 1 % v/v Nonidet P-40, 1 % v/v Triton X-100, 1 % w/v sodium-deoxycholate, 1 mM Na_3_VO_4_, 10 mM NaF, 100 μM leupeptin and 100 μM E64D protease inhibitor) containing cocktails of protease inhibitors and phosphatase inhibitors ^18^. After clarification of the lysates by centrifugation, the total protein concentration was determined by performing a Bradford assay. A standard curve with bovine serum albumin was established.

SDS-PAGE was performed using 10 % w/v polyacrylamide gels to resolve proteins (40 μg per lane) under reducing conditions. Proteins were then transferred by semi-dry transfer onto nitrocellulose membranes. Saturation was performed for 1 h at room temperature in PBS 2 % w/v BSA 0.2 % TWEEN 20 and membranes were incubated overnight at 4 °C with primary antibodies diluted in saturation buffer. For phospho-ERK quantification, combinations of mouse anti-phospho-ERK and rabbit anti-ERK were used. The mCherry-tagged PDE6D was revealed with a mouse anti-PDE6D, and GFP2-tagged K-Ras with a rabbit anti-GFP. Actin staining using a mouse anti-actin antibody was performed as a loading control. Incubation with corresponding secondary antibodies diluted in saturation buffer was carried out for 1 h at room temperature. At least three wash steps in PBS 0.2 % v/v TWEEN 20 were performed after each antibody incubation. An Odyssey Infrared Image System (LI-COR Biosciences) was used to quantify signal intensities. First, the ratio between the intensities obtained for phospho-ERK versus total ERK was calculated for each experimental condition. To make blots comparable by taking into account technical day-to-day variability, normalization to the sum of all the ratios obtained for one blot was done. Finally, data were expressed relative to the averaged control condition and then represented as the mean ± SEM of N = 3 independent biological repeats.

### Quantitative RT-PCR

C2C12 cells were seeded at a density of 100,000 cells/ mL in each well of a 6-well plate and transfected 24 h later with 100 nM of the negative control siRNA or siRNAs targeting mouse *KRAS4B* (NM_001403240.1). Cells were prepared for total RNA extraction after a culture period of 72 h in low serum. NucleoSpin RNA Plus, Mini kit was used to isolate total RNA in accordance with the manufacturer’s instructions. SuperScriptIII Reverse Transcriptase was used to perform reverse transcription on 1 µg of total RNA. SsoAdvanced Universal SYBR Green Supermix on the CFX-connect real-time PCR device (Bio-Rad) and Bio-Rad CFX Manager Software were used to assess the relative abundance of the *KRAS4B* gene transcript. Forward and reverse primer sequences for generating *KRAS* amplicons have been described previously ^30^. The primers for amplification of the *GAPDH* amplicon were created using the online program OligoPerfect Primer Designer. The mRNA sequences of mouse *GAPDH* (NM_008084.4) was used as templates for primer design. The relative mRNA expression level was calculated using the 2-ΔΔCt method by normalizing to GAPDH expression.

### Single-cell RNA sequencing library preparation

C2C12 cells were either cultured in high serum or low serum medium. In addition, siRNA-mediated knock-down of *KRAS4B* or *IFT88* or overexpression of oncogenic K-Ras-G12C were performed under low serum conditions. For single cell analysis, the cells were processed using the Chromium fixed RNA profiling reagent kit for the mouse transcriptome. In brief, the cells were fixed overnight at 4 ℃. After quenching, the cells were hybridized for 17 h with barcoded probes allowing for sample pooling. For post-hybridization washing, the samples with different barcodes were pooled based on the cell count to keep the sample ratio identical in the pool. Pooled samples were loaded on the chip for Gel Beads-in-emulsion (GEM) generation. Prepared libraries were quantified using a Qubit 4 Fluorometer and size distribution was checked on a fragment analyzer. Equimolar pooled libraries were sequenced on NextSeq2000 sequencing system.

### Preprocessing, batch correction, clustering of single-cell RNA sequencing

10x Genomics Cell Ranger (v8.0.1) was used to demultiplex, count and discard empty droplets from raw data of each sample. Correction of ambient RNA was not possible as the complexity of our C2C12 system is low. Filtered count matrices from Cell Ranger were imported and analyzed in R (v 4.3, R Core Team, 2024, https://www.R-project.org/) using Seurat version 5.0.1 ^56, 57^, keeping cells with a minimum of 200 genes and genes with a minimum of 3 cells.

After quality control plots, matrices were filtered to keep cells with the following settings: nCount_RNA < 60000 and percent.mt <= 5. The resulting matrix consisted of 13298 genes across 85776 cells. For clustering and cell type definition, data were normalized using sctransform, regressing out the percentage of mitochondrial genes and reduced by PCA and subsequent by uniform manifold approximation and projection (UMAP) using the first 20 Principal Components. Clustering was then performed using Louvain algorithm for resolutions from 0.1 to 0.8 ^58^. A resolution of 0.3 was chosen for further analysis, based on silhouette and Clustree analysis ^59^. RNA slot of the data were log-normalized and used for further analyses. Cluster markers were found by using FindAllMarkers from Seurat. We looked for positive markers, with the following thresholds: min.diff.pct = 0.1, logfc.threshold = 0.25, min.pct = 0.5. We used MAST as test method ^60^. P-values were corrected using Bonferroni method. For dotplots of single cells, the percentage represented the number of cells expressing the gene, divided by the number of cells in the considered group. The expression is average Seurat log-normalized expression per group, further centered and scaled by gene across groups.

### Pseudotime analysis of single-cell RNA sequencing data

A trajectory was inferred on high serum and low serum C2C12 cell samples representing the unperturbed differentiation using Monocle3 version 1.3.7 ^61^. Briefly, data from Seurat were subset to those samples. Prior to trajectory estimation, clusters were re-estimated on this subset by Monocle using cluster_cells function, and default settings. For the estimation of pseudotime, cluster 6 closest to MuSC.7 cluster was used as a starting point.

### Mathematical models of C2C12 cell differentiation

Two mathematical models were developed in the formalism of ordinary differential equations and fitted to differentiation data of C2C12 cell subpopulations (**Figure S1D**) ^21^. The models have been implemented and analyzed in the IQM toolbox in MATLAB R2019B ^62^. Model states are given in number of cells. Percentages of the different measured cell populations with respect to the overall number of cells were calculated in the models where needed. Time is given in days.

Data integration and estimation of the model parameters were performed within IQM by applying a combined global (*pswamIQM*) and local (*simplexIQM*) optimization. Optimization was repeated 100 times and successful fits with an optimal cost smaller or equal 0.05 were retained for simulation. Due to intrinsic experimental variability the initial conditions were also optimized. Simulation results are shown as the median of all successful model fits (bold dashed line), and as fine lines indicating confidence intervals containing 68 % of the successful fits. Model 1 estimates the fraction of asymmetric cell divisions and employs three states (muscle stem cells with state variable *MuSC*, TAC with the variable *TRANS*, differentiated cells with the variable *DIFF*) and five rate constant parameters. The following assumptions were made for model 1, where (ii) to (iii) are based on our published experimental data ^21^: (i) All rates follow first order kinetics. (ii) The self-renewal rate of MuSC and TAC is reduced to 50% after switching from high to low serum. (iii) The respective rates were directly calculated from the half-life values of cytopainter dye dilution experiments. (iii) Transition from *TRANS* to *DIFF* states occurs only after serum switching. (iv) All cell states are subjected to the same exponential dissipation/ death rate *d*. Model 1 states, parameters and assumptions are shown in **Figure S1A**. The fraction of asymmetric divisions, *xPT*, of the MuSC and TAC was calculated as the sum of the asymmetric rates over the sum of all rates of the *MuSC* and *TRANS* states.

Model 2 integrated ciliation as determinant for asymmetric divisions of ciliated MuSC (variable *MuSCc*) and ciliated TAC (variable *TRANSc*) under differentiating, low serum conditions in addition to states and parameters introduced in model 1. The following assumptions in addition to those from model 1 were made for model 2 captured in: (v) Non-ciliated MuSC and TAC do not asymmetrically divide. (vi) Ciliated MuSC divide into ciliated and non-ciliated MuSC in high serum, while in low serum they divide into ciliated MuSC and non-ciliated TAC. Model 2 states, parameters and assumptions are reflected in **Figure S1C**. The fraction of ciliation in the daughter cells is given by the factor *fRasCil*, a factor ranging from 0 (no ciliation in daughter cells) to 2 (ciliation in both daughter cells). (vii) Ciliated TRANS divide into ciliated and non-ciliated TRANS in high serum and into ciliated transient and non-ciliated differentiated cells in low serum. The fraction of ciliation in the daughter cells is again given by the factor *fRasCil*. The variable *pCil* captures the fraction of ciliated MuSC and TAC cells among all cells.

### Statistical analysis

If not stated otherwise means and standard deviations (SD) are plotted. Data were analyzed using GraphPad Prism 9 or 10 software. The number of independent biological repeats (N) and datapoints (n) for each data set is provided in the figure legends.

Imaging data are representative of phenotypes observed in N ≥ 3 independent biological repeats. Data shown as percent cells, were logit transformed using the cell fraction, p, according to: logit(p)=ln(p/1-p), to obtain continuous values for statistical comparison. Plots were generated such that the Y-axis representing the logit scale was shown on the right and the left Y-axis displayed the approximate percentage values corresponding to the logit scale.

Statistical tests are specified in the figure legends. Comparisons were done against controls (typically plotted in grey) or as indicated. A p-value of < 0.05 was considered statistically significant and the statistical significance levels were annotated as: * = P < 0.05; ** = P < 0.01; *** = P < 0.001; **** = P < 0.0001, or ns = not significant.

## Acknowledgements

We thank Lynn Mangen, Nesrine Ben Fredj and Dr. Farah Kouzi for their experimental support. Our thanks go to Olga Kondratyeva and Dr. Paul Antony (bioimaging platform, LCSB, University of Luxembourg) for their assistance with the set-up of FACS-sorting. We also thank Dr. Rashi Halder (genomics platform, LCSB, University of Luxembourg) for running scRNAseq samples. We are thankful to Gesa Hoelzer and Anna Kemeter of the Jena lab for supporting the FRAP measurements and providing a python code for analysis of FRAP data. We thank the Microverse Imaging Center Jena and Aurélie Jost / Patrick Then for providing microscope facility support for data acquisition and data analysis.

## Author contributions

D.K.A conceived the project and supervised the study. R.C., E.S.R., B.P., S.B., C.L., Y.R., M.D., and A.G.M. conducted experiments, analyzed and interpreted data and prepared figures. A.G. evaluated scRNAseq data. T.S. performed the mathematical modelling. J.d.H. supervised zebrafish experiments and C.E. oversaw super-resolution STED-microscopy experiments. D.K.A., J.d.H., and C.E. acquired funding. R.C., E.S.R. contributed to the drafting and D.K.A. wrote the final manuscript. All authors gave final approval and agreed to be accountable for all aspects of work, ensuring integrity and accuracy.

## Ethical approval

All procedures involving experimental animals are approved by the local animal experiments committee (AVD8010020173786; license valid until 14 December 2027).

## Data and Code Availability

The mouse C2C12 cell line single-cell RNA sequencing data described here are available at the ENA database under the accession reference PRJEB89780. Code used in this study is listed in the Key Resources Table and has been deposited in Gitlab at https://gitlab.com/uniluxembourg/fstm/dlsm/ccbdd/chippalkatti_2025, GitHub at https://github.com/sysbiolux/RasCilium and https://github.com/GesaHlzer/BachelorThesis.

## Additional information

### Supplementary information

The online version contains supplementary material available at [link]

### Correspondence and requests for materials

Further information and requests for resources and reagents should be directed to and will be fulfilled by the lead contact, Prof. Dr. Daniel Kwaku Abankwa.

