## Supplementary Information for "K-Ras controls asymmetric cell divisions from the primary cilium"

#### Supplementary Figures

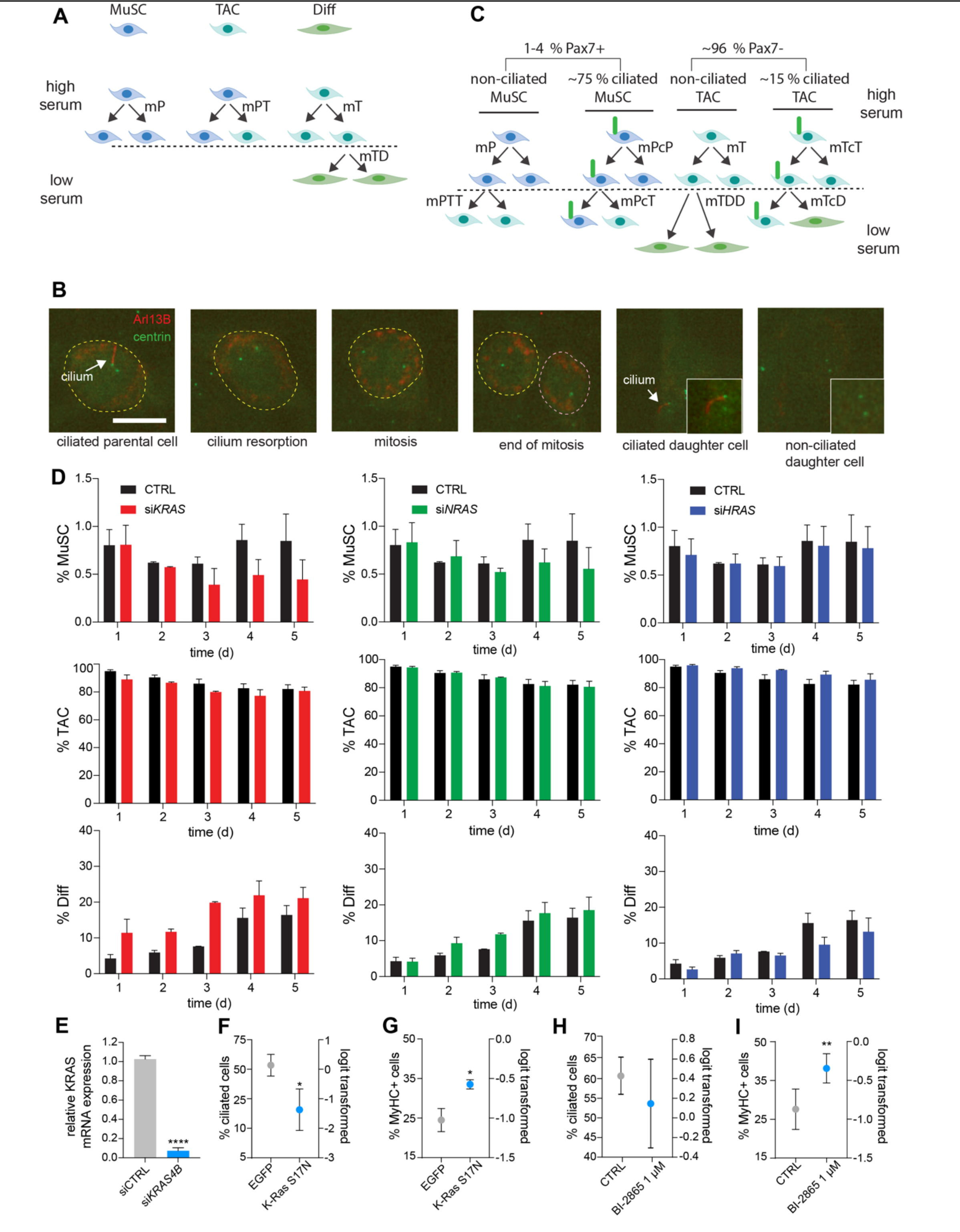


**Figure S1. Related to Figure 1.**

(**A**) Schematic of mouse skeletal muscle C2C12 cell hierarchy with rate constants used for the first ordinary differential equation model (model 1). The decay/ death rate d is not displayed, as it impacts equally on all states.

This simple model could adequately describe the evolution of MuSC and TAC in high serum, where cells proliferate. It likewise provided a good fit to data from low serum conditions, where a very low percentage of MuSC and a high percentage of TAC decrease, while differentiated cells (Diff) increase.

(**B**) Confocal images of time-lapse recording of a C2C12 cell grown in high serum for 48 h after transfection with Arl13B-mCherry as ciliary marker (arrow) and EGFP-centrin1 to mark the centrioles. Scale bar = 10 μm. The corresponding video is provided as **Video S1**.

(**C**) Schematic of C2C12 cell hierarchy with rate constants used for the second ordinary differential equation model (model 2), which included ciliation as determinant for asymmetric cell divisions. The decay/ death rate d is not displayed, as it impacts equally on all states.

(**D**) Flow-cytometric quantification of C2C12 cell sub-populations, MuSC (Pax7+/MyHC-), TAC (Pax7-/MyHC-) and Diff cells (Pax7-/MyHC+) after induction of differentiation in low serum. Indicated knockdowns of Ras isoforms were done on day 0 using 100 nM siRNAs, N = 3. Means ± SD are plotted. The time-aggregated data were published by us before [1].

(**E**) RT-PCR based *KRAS* mRNA levels normalized to *GAPDH* mRNA of C2C12 cells grown in low serum for 72 h after transfection with siCTRL or si*KRAS4B* (each 100 nM), N = 4. Means ± SD are plotted. Statistical analysis was done with the unpaired t-test.

(**F**) Confocal imaging-based quantification of ciliation of C2C12 cells grown in high serum for 48 h after transfection with EGFP alone or EGFP-K-RasS17N, N = 3, n ≥ 250. Means ± SD are plotted. Statistical analysis was done with the unpaired t-test.

(**G**) Flow cytometric quantification of MyHC terminal differentiation marker expression of C2C12 grown in low serum for 72 h and transfected as in (F), N = 3. Means ± SD are plotted. Statistical analysis was done with the unpaired t-test.

(**H**) Confocal imaging-based quantification of ciliation of C2C12 grown in low serum for 72 h and treated with CTRL (DMSO 0.1 % in low serum medium) or pan-K-Ras inhibitor BI-2865 at the indicated concentration, N = 3, n ≥ 250. Means ± SD are plotted. Statistical analysis was done with the unpaired t-test.

(**I**) Flow cytometric quantification of MyHC terminal differentiation marker expression in C2C12 treated as in (H), N = 3. Means ± SD are plotted. Statistical analysis was done with the unpaired t-test.

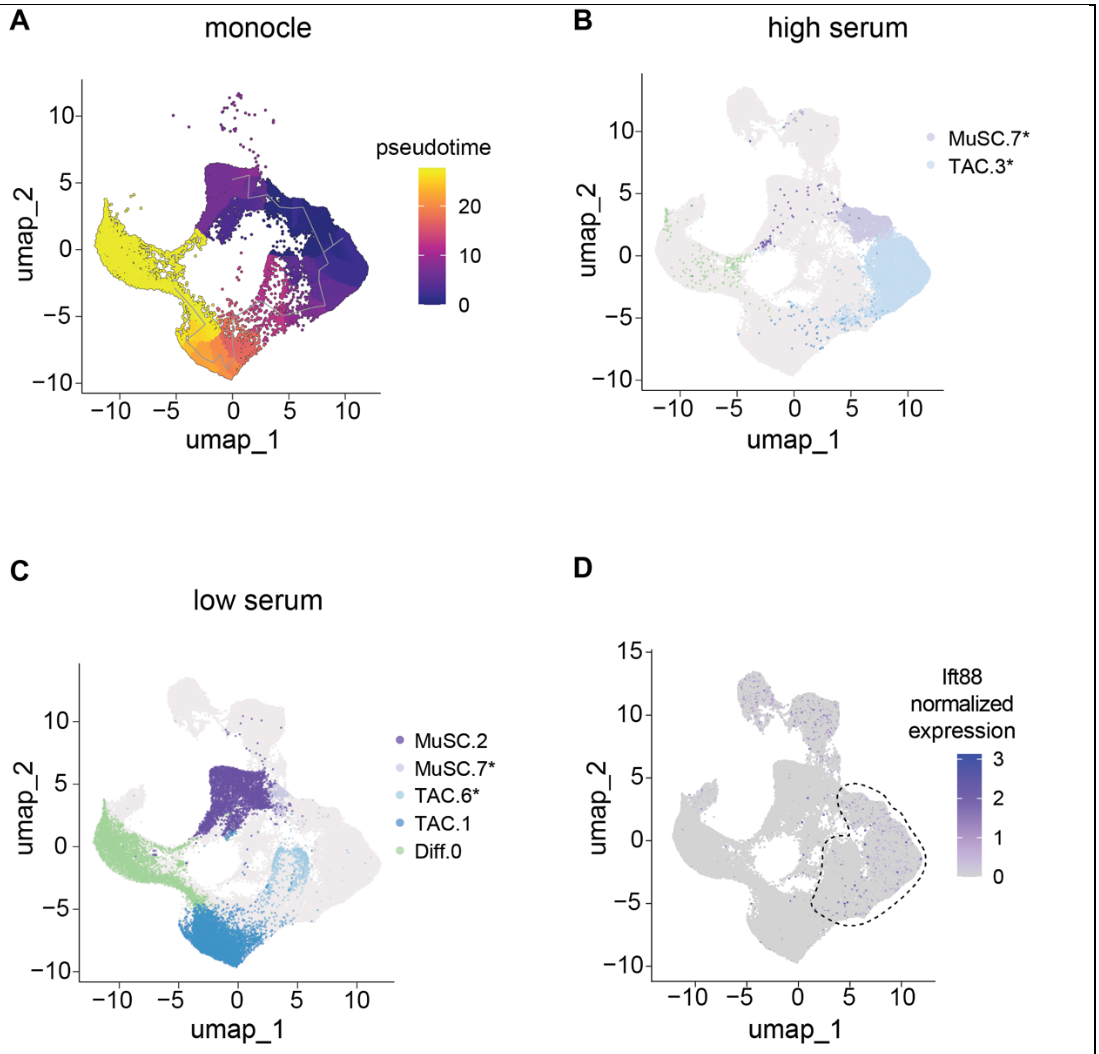


**Figure S2. Related to Figure 2.**

(**A**) Prediction of mouse skeletal muscle C2C12 cell differentiation trajectory and pseudotime with Monocle.

(**B, C**) UMAP projection of C2C12 cell scRNAseq data, with cells from high serum (B) and from samples of cells cultured 72 h in low serum (C) colored by cluster. Bona fide ciliated cell clusters MuSC.7, TAC.3 and TAC.6 are marked with an asterisk.

(**D**) Ift88-expression overlaid on UMAP projection of C2C12 cell scRNAseq data. Ciliated cell clusters MuSC.7, TAC.3 and TAC.6 are outlined with a dashed line.

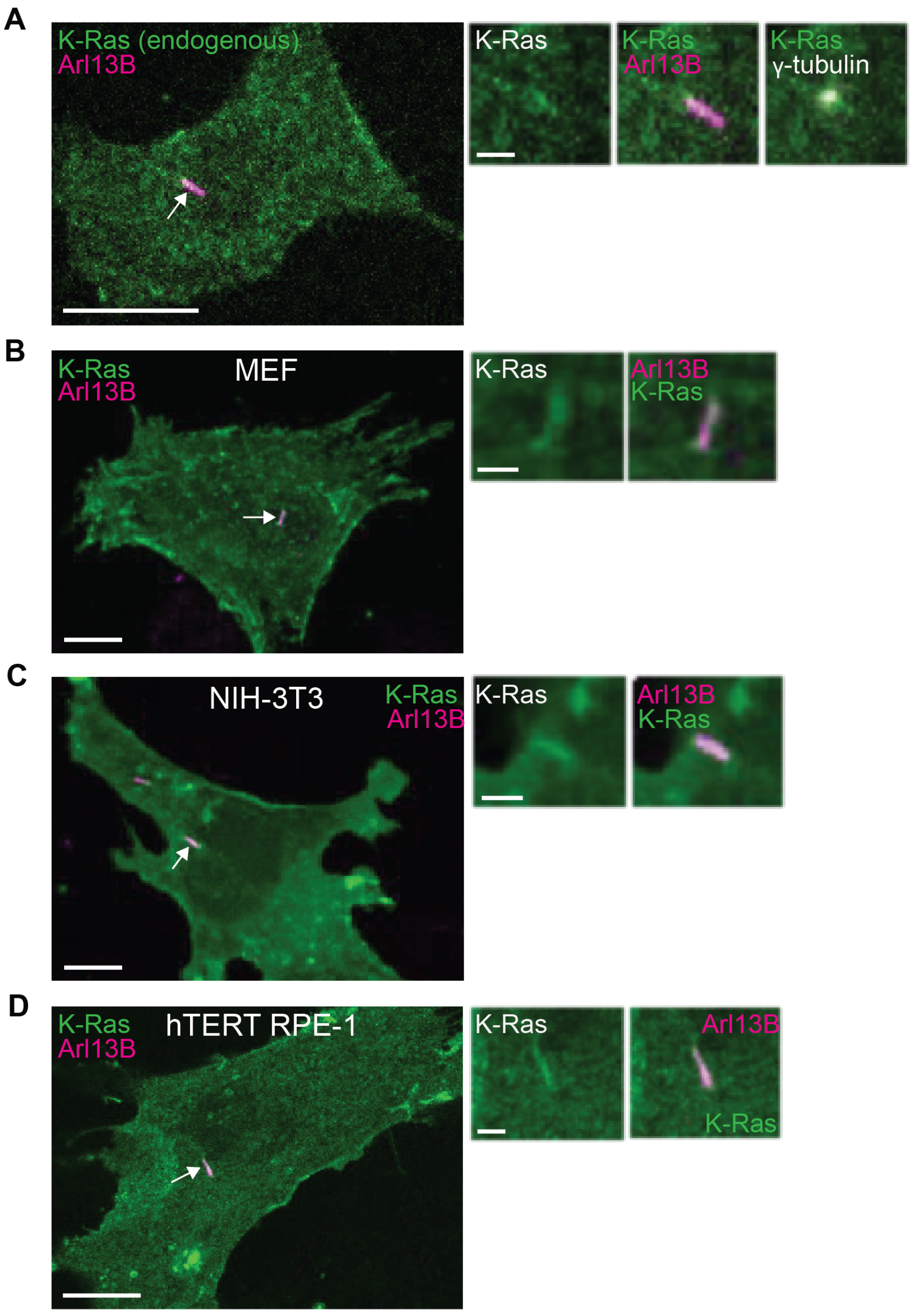


**Figure S3. Related to Figure 3.**

(**A**) Confocal image of mEos3.2-KRAS knock-in C2C12 cells cultured in high serum for 24 h directly after FACS-sorting and immunolabelled for ciliary marker Arl13B (arrow) and centriolar marker γ-tubulin. Scale bar = 10 µm. Smaller images to the right show enlargement of the area containing the cilium. Scale bar = 2 µm.

(**B-D**) Confocal images of MEF wt (B), NIH-3T3 (C) and hTERT RPE-1 cells (D) that were transfected with mEGFP-K-Ras. 24 h after transfection, cells were serum starved for 24 h to induce the formation of a primary cilium and immunolabelled for ciliary marker Arl13B (arrow). Scale bar = 10 µm. Smaller images to the right show enlargement of the area containing the cilium. Scale bar = 2 µm.

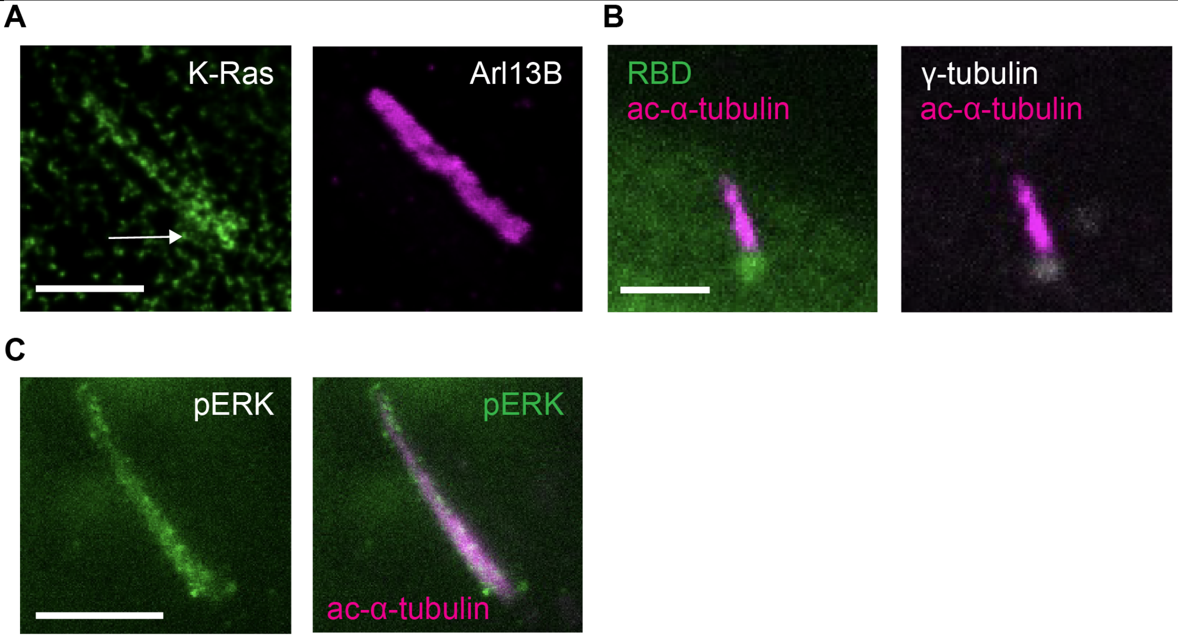


**Figure S4. Related to Figure 4.**

(**A**) STED images of C2C12 cells grown in high serum for 48 h after transfection with GFP2-K-Ras and immunolabeled for GFP with ATTO647N-conjugated nanobody and ciliary membrane marker Arl13B. At the base of the cilium, K-Ras appears to be distributed in a radially symmetrical fashion (arrow), as would be expected from its association with the centrioles. Scale bar = 2 μm.

(**B**) Confocal images of C2C12 cells grown in high serum for 48 h after transfection with mCherry-C-Raf-RBD and immunolabelled for acetylated α-tubulin marking the cilium. Cells were not transfected with K-Ras variant constructs, thus showing dependence on endogenous active Ras. Scale bar = 2 μm.

(**C**) Confocal images of C2C12 cells grown in high serum for 48 h. Cells were not co-transfected with K-Ras variant constructs. Immunolabelling for endogenous phospho-Thr202/Tyr204-ERK1/2 and acetylated α-tubulin. Scale bar = 2 μm.

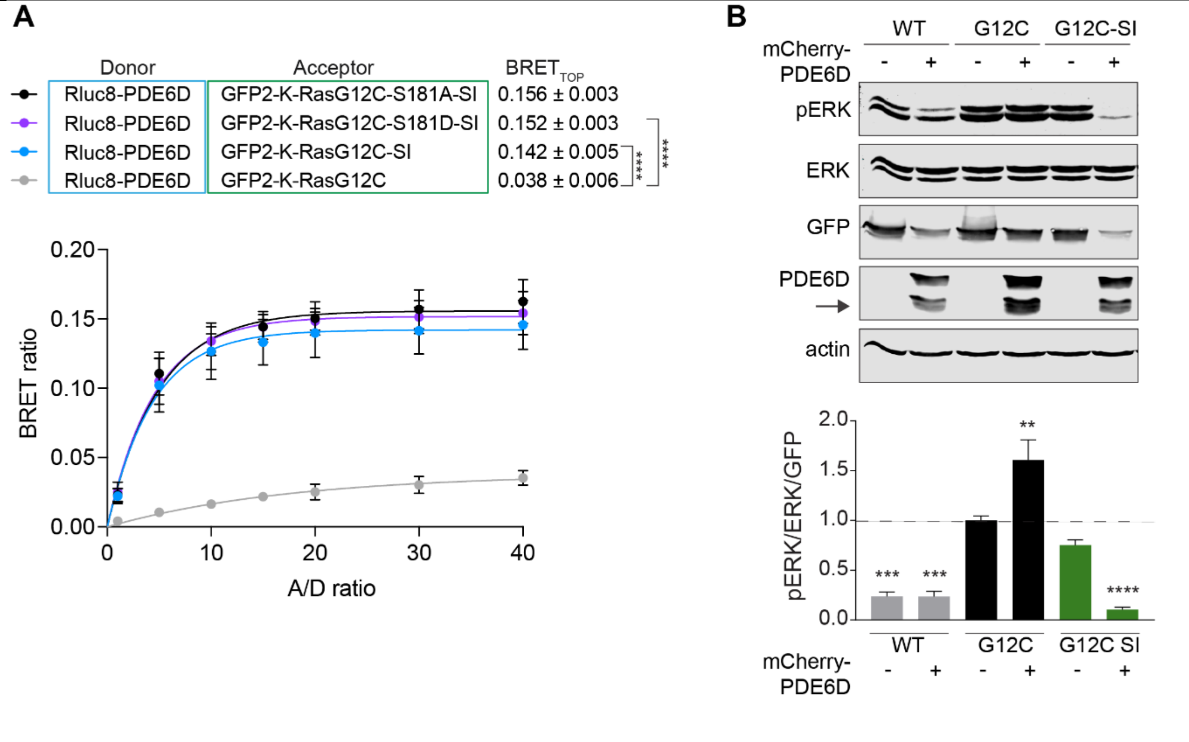


**Figure S5. Related to Figure 5.**

(**A**) BRET titration curves of Rluc8-PDE6D and GFP2-K-RasG12C mutants expressed in HEK cells, N = 3. Statistical analysis was performed with the extra sum-of-squares F-test.

(**B**) Immunoblot data (top) with quantified normalized phosphorylated ERK (pERK) levels after expression of EGFP-K-Ras (WT), EGFP-K-RasG12C (G12C) or EGFP-K-RasG12C-SI (G12C-SI) expressed in HEK cells with or without co-expression of mCherry-PDE6D at 1:1 DNA ratio for 24 h, N = 3. Means ± SEM are plotted. Employed antibody labelings are indicated to the left of the blot. Statistical analysis was done using One-way-ANOVA with multiple comparisons with G12C without mCherry-PDE6D expression used as a reference. Arrow marks truncated/ cleaved protein consistently observed following transfection of mCherry-PDE6D.

#### Video S1

C2C12 cells were transfected with Arl13B-mCherry and mEGFP-centrin. A cilium as well as the mother and daughter centrioles are clearly observed at t = 00:00:00 in the cell marked with a circle. The cilium is resorbed at t = 01:00:00 and the cell enters mitosis at approximately t = 02:00:00. The two daughter cells subsequently generated are marked with yellow and pink circles at t = 02:50:00. Both daughter cells then traverse for a short distance and only one, indicated with a pink border assembles a cilium at t = 08:50:00. This indicates asymmetric inheritance of the cilium in C2C12 cell divisions. Images were acquired at a time interval of 10 min and 29 z-stacks were acquired with a spacing of 0.5 µm. Time-lapse movie represents z-projection at maximum intensity.

**Table S1: Materials and equipment employed in the study**

| REAGENT or RESOURCE | SOURCE | IDENTIFIER |
|---|---|---|
| Antibodies | | |
| Mouse monoclonal anti-Pax7 (Pax7);<br>Dilution 1:500 immunofluorescence (IF) | Bio-Techne | Cat# MAB1675<br>RRID:<br>AB_2159833 |
| Rabbit polyclonal anti-Arl13B;<br>Dilution 1:500 IF | Proteintech | Cat# 17711-1-AP<br>RRID:<br>AB_2060867) |
| Alexa Fluor 647 Phalloidin;<br>Dilution 1:50 IF | Invitrogen | Cat# A-22287<br>RRID:<br>AB_2620155 |
| ChromoTek GFP-Booster ATTO647N;<br>Dilution 1:200 IF | Proteintech | Cat# gba647n<br>RRID:<br>AB_2629215 |
| Mouse monoclonal anti-alpha tubulin (acetyl K40) [EPR16772];<br>Dilution 1:500 IF | Abcam | Cat# ab179484<br>RRID:<br>AB_2890906 |
| Rabbit recombinant monoclonal anti-gamma-2- tubulin antibody [EPR16793];<br>Dilution 1:500 IF | Abcam | Cat# ab179503<br>RRID:<br>AB_2904198 |
| Rabbit polyclonal anti-Phospho MEK1/2 (Ser217/221);<br>Dilution 1:200 IF | Cell Signaling | Cat# 9121<br>RRID: AB_331648 |
| Rabbit polyclonal anti-Phospho-p44/42 MAPK (Erk1/2) (Thr202/Tyr204) Antibody<br>Dilution 1:200 IF | Cell Signaling | Cat# 9101<br>RRID: AB_331646 |
| Goat anti-Rabbit IgG (H+L) cross-adsorbed secondary antibody, Alexa Fluor 594;<br>Dilution 1:200 IF | ThermoFisher Scientific | Cat# A-11012<br>RRID:<br>AB_2534079 |
| Goat anti-mouse IgG (H+L) cross-adsorbed secondary antibody, Alexa Fluor 647;<br>Dilution 1:200 IF | ThermoFisher Scientific | Cat# -21235<br>RRID:<br>AB_2535804 |

| | | |
|---|---|---|
| Anti-Rabbit IgG - Atto 633 antibody produced in goat<br>Dilution 1:200 IF | Merck | Cat# 41176<br><br>RRID:<br>AB_1137641 |
| Mouse monoclonal anti-myosin 4 antibody (MF20), eFluor660, eBioscience;<br>Dilution 1:100 Flow cytometry | ThermoFisher Scientific | Cat# -50-6503-80<br>RRID:<br>AB_2574266 |
| Rabbit polyclonal anti-p44/42 MAPK (Erk1/2);<br>Dilution 1:1,000 Western Blot (WB) | Cell Signaling | Cat# 9102<br>RRID: AB_330744 |
| Mouse monoclonal anti-phospho-p44/42 MAPK (Erk1/2) (Thr202/Tyr 204) (E10);<br>Dilution 1:2,000 WB | Cell Signaling | Cat# 9106<br>RRID: AB_331768 |
| Mouse monoclonal anti-PDE6D (E7);<br>Dilution 1:1,000 WB | Santa Cruz Biotechnology | Cat# sc-166854<br>RRID:<br>AB_2161460 |
| Rabbit polyclonal anti-GFP antibody;<br>Dilution 1:5,000 WB | Merck | Cat# SAB4301138<br>RRID:<br>AB_2750576 |
| Mouse monoclonal anti-beta actin antibody;<br>Dilution 1:5,000 WB | Merck | Cat# A5441<br>RRID: AB_476744 |
| IRDye 680RD goat anti-Rabbit IgG;<br>Dilution 1:10,000 WB | LI-COR Biosciences | Cat# 926-68071<br>RRID:<br>AB_10956166 |
| IRDye 680LT donkey anti-Mouse IgG;<br>Dilution 1:10,000 WB | LI-COR Biosciences | Cat# 926-68022<br>RRID:<br>AB_10715072 |
| IRDye 800CW goat anti-Rabbit IgG;<br>Dilution 1:10,000 WB | LI-COR Biosciences | Cat# 926-32211<br>RRID: AB_621843 |
| IRDye 800CW donkey anti-Mouse IgG;<br>Dilution 1:10,000 WB | LI-COR Biosciences | Cat# 926-32212<br>RRID: AB_621847 |
| Bacterial and virus strains | | |
| *E. coli* DH10B | New England BioLabs | Cat# C3019I |
| Biological samples | | |
| N/A | N/A | N/A |

| | | |
|---|---|---|
| Chemicals, peptides, and recombinant proteins | | |
| BI-2865 | Selleckchem | Cat# E1474 |
| Dimethyl sulfoxide (DMSO) | PanReac AppliChem | Cat# A3672 |
| Hoechst 33342 | ThermoFisher Scientific | Cat# 62249 |
| Critical commercial reagents | | |
| Gateway LR Clonase II enzyme mix | ThermoFisher Scientific | Cat# 11791020 |
| Gateway BP Clonase II enzyme mix | ThermoFisher Scientific | Cat# 11789020 |
| Bio-Rad Protein Assay Kit II | Bio-Rad | Cat# 5000002 |
| Pierce Protease Inhibitor Tablets | ThermoFisher Scientific | Cat# A32963 |
| PhosSTOP | Merck | Cat# 4906845001 |
| Lipofectamine RNAiMAX Transfection Reagent | ThermoFisher Scientific | Cat# 13778075 |
| Lipofectamine 2000 Transfection Reagent | ThermoFisher Scientific | Cat# 11668019 |
| jetPRIME transfection reagent | Polyplus | Cat#101000046 |
| Coelenterazine 400a; 2,8-Dibenzyl-6-phenylimidazo[1,2a]pyrazin-3-(7H)-one; DeepBlueC | Gold Biotechnology | Cat# C-320-1 |
| NucleoSpin RNA Plus, Mini kit for RNA purification | Macherey-Nagel | Cat# 740984.250 |
| SuperScriptIII Reverse Transcriptase | ThermoFisher Scientific | Cat# 18080093 |
| SsoAdvanced Universal SYBR Green Supermix | Bio-Rad | Cat# 1725274 |
| Rapid DNA ligation kit | ThermoFisher Scientific | Cat# K1422 |
| Shrimp Alkaline Phosphatase (rSAP) | New England Biolabs | Cat# M0371S |
| BsrGI-HF | New England Biolabs | Cat# R3575S |
| NheI-HF | New England Biolabs | Cat# R3131S |
| NotI | New England Biolabs | Cat# R0189S |
| SpeI-HF | New England Biolabs | Cat# R3133S |
| PstI | New England Biolabs | Cat# R0140S |
| XbaI | New England Biolabs | Cat# R0145S |
| SalI | New England Biolabs | Cat# R0138S |
| NEBuffer r2.1 | New England Biolabs | Cat# B6002S |
| NEBuffer r3.1 | New England Biolabs | Cat# B6003S |
| rCutSmart Buffer | New England Biolabs | Cat# B6004S |

| | | |
|---|---|---|
| Q5 High fidelity DNA polymerase | New England Biolabs | Cat# M0491L |
| dNTP set, 100 mM | ThermoFisher Scientific | Cat# R0181 |
| mMessage mMachine SP6 transcription kit | ThermoFisher Scientific | Cat# AM1340 |
| Agarose | Merck | Cat# 2120 |
| NucleoSpin Gel and PCR clean-up kit, mini kit for gel extraction or PCR clean-up | Macherey-Nagel | Cat# 740609.50 |
| Experimental models: Cell lines | | |
| Mouse cell line, C2C12 | ATCC | Cat# CRL-1772 RRID: CVCL_0188 |
| Mouse cell line, C2C12-mEos3.2-KRAS knock-in | Synthego | N/A |
| Human cell line, HEK293-EBNA (HEK) | Prof. Florian M. Wurm, EPFL | RRID: CVCL_6974 |
| Human cell line, 293 c18 | ATCC | Cat# CRL-10852, RRID: CVCL_6974 |
| Mouse cell line, wt MEFs | ATCC | Cat# CRL-2991, RRID: CVCL_L69 0 |
| Mouse cell line, NIH-3T3 | ATCC | Cat# CRL-1658 RRID: CVCL_0594 |
| Human cell line, hTERT RPE-1 | ATCC | Cat# CRL-4000 RRID:CVCL_4388 |
| Experimental models: Organisms/strains | | |
| N/A | N/A | N/A |
| Oligonucleotides | | |
| Custom qPCR primer: KRAS mouse_F 5’-3’sequence GGAGTACAGTGCAATGAGGGAC | Genecust | N/A |
| Custom qPCR primer: KRAS mouse_R 5’-3’sequence CCAGGACCATAGGCACATCTTC | Genecust | N/A |
| Custom qPCR primer: GAPDH mouse_F 5’-3’ sequence TGGTGAAGGTCGGTGTGAA | Eurogentec | N/A |
| Custom qPCR primer: GAPDH mouse_R | Eurogentec | N/A |

| | | |
|---|---|---|
| 5'-3' sequence ATGAAGGGGTCGTTGATGG | | |
| Custom Restriction cloning primer: PstI KRAS S17N Forward<br>5'-3' sequence: TCACTGCAGGCATGACTGAATATAAA CTTGTGGTAGTTG | Integrated DNA technologies | N/A |
| Custom Restriction cloning primer: SpeI KRAS S17N Reverse<br>5'-3' sequence: CTGACTAGTTTACATAATTACACACTT TGTCTTTGACT | Integrated DNA technologies | N/A |
| Custom Restriction cloning primer: NotI KRAS SI Forward<br>5'-3' sequence: GTGGCGGCCGCATGGTGAGCAAGGG CG | Integrated DNA technologies | N/A |
| Custom Restriction cloning primer: SpeI KRAS SI Reverse<br>5'-3' sequence: ACTACTAGTTCAAGAAACGGAGCAGA TGG | Integrated DNA technologies | N/A |
| Human Negative control siRNA | QIAGEN | Cat# 1027310 |
| Human *HRAS* siRNA (FlexiTube siRNA)<br>5'-3'sequence<br>CGGAAGCAGGUGGUCAUUA | QIAGEN | Cat# 1027417 |
| Mouse *KRAS* siRNA (targeting both *KRAS*4A and 4B transcripts; si-pan *KRAS*)<br>(ON-TARGETplus SMARTpool siRNA)<br>5'-3' sequences<br>GAACAGUAGACACGAAAA<br>AGCAAGGAGUUACGGGAUU<br>GGUUGGAGCUGGUGGCGUA<br>GGUGUACAGUUAUGUGAAU | Horizon Discovery | Cat# L-043846-01-0005 |

| | | |
|---|---|---|
| Custom siRNA: Mouse *NRAS* siRNA 5'-3' sequence GCAAAUUAAGCGUGUGAAAUUU | Horizon Discovery | N/A |
| Custom siRNA: Mouse *KRAS4B* siRNA 5'-3' sequence Cy3-AAGAAGAAGUCAAGGACAAUU | Horizon Discovery | N/A |
| Mouse *PDE6D* siRNA (ON-TARGETplus SMARTpool siRNA) 5'-3' sequences UCUCAUACCUUUAACUUGU CGCCUGAGUCCCAGAUGAU CAAAGGACAAUGCCUAGAA GCGCAAACAUGGGAACAAA | Horizon Discovery | Cat# L-062279-01-0005 |
| Recombinant DNA | | |
| C453-E04: attB4-CMV51p>-attB5 (L4 L5) | FNL Combinatorial Cloning Platform, RAS-Initiative | Addgene# 162973 |
| C413-E36: attB4-CMV51p>-attB1r (L4 R1) | FNL Combinatorial Cloning Platform, RAS-Initiative | Addgene# 162927 |
| pDest-305 (attR4-attR2) | FNL Combinatorial Cloning Platform, RAS-Initiative | Addgene# 161895 |
| pDest-312 (attR4-attR3) | FNL Combinatorial Cloning Platform, RAS-Initiative | Addgene# 161897 |
| C512-E01: attB5r-meGFP-nostop-attB1r (R5 R1) | FNL Combinatorial Cloning Platform, RAS-Initiative | Addgene# 162940 |
| C511-E03: attB5r-Rluc8-nostop-attB1r (R5 R1) | [2] | N/A |
| pDONR235 GFP2-no stop R5 R1 | Generated by Genewiz Inc. | N/A |

| | | |
|---|---|---|
| C512-E04: attB5r-mCherry-nostop-attB1r (R5 R1) | FNL Combinatorial Cloning Platform, RAS-Initiative | Addgene# 162941 |
| C232-E06: attB2r-mCherry-attB3 (R2 L3) | FNL Combinatorial Cloning Platform, RAS-Initiative | Addgene# 162903 |
| Gateway pDONR221 vector | Thermofisher Scientific | Cat# 12536017 |
| Hs. PDE6D | [2] | N/A |
| Hs. KRAS4B | RAS mutant collection V2.0, RAS-Initiative | Addgene# 83129 |
| Hs. HRAS WT | RAS mutant collection V2.0, RAS-Initiative | Addgene# 83181 |
| Hs. KRAS4B-S181A | RAS mutant collection V2.0, RAS-Initiative | Addgene# 83136 |
| Hs. KRAS4B-S181D | RAS mutant collection V2.0, RAS-Initiative | Addgene# 83137 |
| Hs. BRAF | RAS pathway clone collection V2.0, RAS-Initiative | Addgene# 70299 |
| pDest305-CMV-GFP2-K-Ras4B | This paper | N/A |
| pDest305-CMV- GFP2-H-Ras | This paper | N/A |
| pDest305-CMV- GFP2-N-Ras | This paper | N/A |
| pDest305-CMV- Rluc8-PDE6D | This paper | N/A |
| pDest305-CMV- mCherry-PDE6D | This paper | N/A |
| pDest305-CMV- GFP2-B-Raf | This paper | N/A |
| pDest305-CMV- mCherry-B-Raf | This paper | N/A |
| pmEGFP-N-Ras | [3] | N/A |
| pDest305-CMV-mEGFP-K-Ras4B | [1] | |
| pDest305-CMV-mEGFP-K-Ras4B G12C | [1] | |
| K-Ras-SI g-block | Integrated DNA technologies | N/A |
| pDest305-CMV-GFP2-K-Ras4B-SI | This paper | N/A |
| pmEGFP-K-Ras SI | Generated by Genecust | N/A |
| pDest305-CMV-GFP2-K-Ras4B G12C-SI | Generated by Genecust | N/A |
| pDest305-CMV-GFP2-K-Ras4B G12C S181A-SI | Generated by Genecust | N/A |

| | | |
|---|---|---|
| pDest305-CMV-GFP2-K-Ras4B G12C S181D-SI | Generated by Genecust | N/A |
| pEGFP-C1 | NovoPro | Cat# V012024 |
| pcDNA3.1(+)mCherry-K-Ras4B S17N | [4] | N/A |
| pEGFP-K-Ras4B S17N | This paper | N/A |
| pCS2+-CMV-mEGFP-K-Ras4B-wt | This paper | N/A |
| pCS2+-CMV-mEGFP-K-Ras4B-SI | This paper | N/A |
| pCS2+-CMV-mEGFP-K-Ras4B-S17N | This paper | N/A |
| pcDNA3.1(+) | Invitrogen | Cat# V79020 |
| pmRFP1-C-RBD | [3] | N/A |
| pEGFP-C1-Centrin | Addgene | Cat# 72641 |
| Mouse Arl13B codon optimized g-block | Integrated DNA technologies | N/A |
| pDest305-CMV-Arl13B-mCherry | This paper | N/A |
| Software and algorithms | | |
| GuavaSoft version4.0 | Cytek Biosciences | N/A |
| Leica Application Suite X version 4.12 | Leica Microsystems | RRID: SCR_013673 |
| GraphPad Prism version 9.5.1 | GraphPad | RRID: SCR_002798 |
| Fiji version 2.9.0 | [5] | RRID: SCR_002285 |
| FlowFate version 1.2 | [6] | N/A |
| Image Studio software version 5.2 | LI-COR Biosciences | N/A |
| Bio-Rad CFX Manager version 3.1 | Bio-Rad | N/A |
| FlowJo version 10.8.2 | FlowJo LLC | RRID:SCR_008520 |
| OriginPro version 2021 | OriginLab | RRID:SCR_014212 |
| SnapGene version 8.0.2 | SnapGene | RRID:SCR_015052 |
| Cell Ranger version 8.0.1 | 10x Genomics | RRID:SCR_017344 |
| R version 4.3 | R Core Team | RRID: SCR_003005 |

| | | |
|---|---|---|
| Seurat version 5.0.1 | 7 | RRID:SCR_016341 |
| Monocle3 version 1.3.7 | 8 | RRID:SCR_018685 |
| MAST | 9 | RRID:SCR_016340 |
| Clustree | 10 | RRID:SCR_016293 |
| Louvain algorithm | 11 | N/A |
| Python version 3.13.3 | Python software foundation | https://www.python.org/ RRID:SCR_008394 |
| Fitting-curves-python: curve fitting algorithm | This paper | https://github.com/GesaHlzer/BachelorThesis |
| **Other** | | |
| Guava easyCyte 6HT 2L flow cytometer | Cytek Biosciences | Cat# 0500-4007 |
| Odyssey CLx Infrared Imaging System | LI-COR Biosciences | Cat# LI9140-00 |
| CFX-connect real-time PCR Detection System | Bio-Rad | Cat# 1855200 |
| 4D-Nucleofector® X Unit | Lonza | Cat# AAF-1003X |
| Qubit 4 Fluorometer | ThermoFisher Scientific | Cat# Q33238 |
| NextSeq2000 sequencing system | Illumina | Cat# 20038897 |
| 5200 Fragment Analyzer 12 capillary electrophoresis system | Agilent | Cat# M5310AA |
| SE Cell Line 4D-Nucleofector™ X Kit L | Lonza | Cat# V4XC-1024 |
| Chromium fixed RNA profiling reagent kit, mouse transcriptome | 10x Genomics | Cat# PN-1000496 |
| dsDNA Quantitation kit, HS | ThermoFisher Scientific | Cat# Q32854 |
| NextSeq™ 2000 P4 XLEAP-SBS™ Reagent Kit | Illumina | Cat# 20100994 |
| Dulbecco's Modified Eagle's Medium (DMEM), high glucose | ThermoFisher Scientific | Cat# 11965092 |

| | | |
|---|---|---|
| Dulbecco's Modified Eagle Medium/Nutrient Mixture F-12 (DMEM/F-12) | ThermoFisher Scientific | Cat# 11320033 |
| DMEM, high glucose, no glutamine, no phenol red | ThermoFisher Scientific | Cat# 31053028 |
| Fetal bovine serum (FBS) | ThermoFisher Scientific | Cat# 10270106 |
| Horse serum (HS) | ThermoFisher Scientific | Cat# 16050130 |
| L-glutamine (200 mM) | ThermoFisher Scientific | Cat# A2916801 |
| Penicillin-Streptomycin (10,000 U/mL) | ThermoFisher Scientific | Cat# 10378016 |
| Hygromycin B (50mg/mL in PBS) | Takara Bio | Cat# 631309 |
| Opti-MEM Reduced Serum Medium | ThermoFisher Scientific | Cat# 31985047 |
| Paraformaldehyde 16 % (w/v) | Avantor | Cat# 30525-89-4 |
| Bovine Serum Albumin | AppliChem | Cat# A6588 |
| TWEEN 20 | Merck | Cat# P9416 |
| Triton X-100 | Merck | Cat# T8787 |
| Semi-dry transfer kit | Bio-Rad | Cat # 1704272 |
| VECTASHIELD Anti-fade mounting medium | Vector Laboratories | Cat# H-1000-10 |
| MOWIOL 4-88 reagent | Merck | Cat# 475904 |
| Cellview cell culture dish, polystyrene, 35/10 mm, glass bottom | Greiner | Cat# 627860 |
| VWR 6-well cell culture plate | Avantor | Cat# 10062-892 |

#### Supplementary Code

MATLAB code for modelling of C2C12 cell differentiation. All rate constants are graphically explained in **Figure S1A,C**.

**Model1 in IQM toolbox format:**

```
********** MODEL NAME
RASdiff

********** MODEL NOTES
RASdiff, T. Sauter, University of Luxembourg, 01-09/24

********** MODEL STATES
d/dt(MuSC) = r_mP - r_dMuSC
d/dt(TRANS) = r_mPT + r_mT - r_mTD - r_dTRANS
d/dt(DIFF) = 2 * r_mTD - r_dDIFF

MuSC(0) = 981
TRANS(0) = 10249
DIFF(0) = 0

********** MODEL PARAMETERS
sP_PT = 1.16
mT = 1.36
xPT = 1
mTD = 1
d = 1
serumSwitch = 0

********** MODEL VARIABLES
serum = 1 - serumSwitch
PAX7neg = TRANS + DIFF
sumPTD = MuSC + TRANS + DIFF
pMuSC = MuSC/sumPTD * 100
pTRANS = TRANS/sumPTD * 100
pDIFF = DIFF/sumPTD * 100

********** MODEL REACTIONS
```

r_mP = sP_PT*(1-xPT) * MuSC *(1+serum)

r_mPT = sP_PT*xPT * MuSC *(1+serum)

r_mT = mT * TRANS *(1+serum)

r_mTD = serumSwitch * mTD * TRANS

r_dMuSC = d * MuSC

r_dTRANS = d * TRANS

r_dDIFF = d * DIFF

********** MODEL FUNCTIONS

********** MODEL EVENTS

********** MODEL MATLAB FUNCTIONS

===

**Model2 in IQM toolbox format:**

********** MODEL NAME

RASdiff

********** MODEL NOTES

RASdiff, T. Sauter, University of Luxembourg, 01-10/24

********** MODEL STATES

d/dt(MuSC) = r_mP - r_mPTT - r_dMuSC + r_mPcP - rasDiff*r_mPcP*(fRasCil-1) + rasDiff*(1-min(1,fRasCil))*mPcT

d/dt(TRANS) = r_mT + 2*r_mPTT - r_mTDD - r_dTRANS + r_mPcT + r_mTcT - rasDiff*r_mTcT*(fRasCil-1) + rasDiff*(1-min(1,fRasCil))*mTcD

d/dt(DIFF) = 2*r_mTDD - r_dDIFF + r_mTcD

d/dt(MuSCc) = - r_dMuSCc + rasDiff*r_mPcP*(fRasCil-1) - rasDiff*(1-min(1,fRasCil))*mPcT

d/dt(TRANSc) = - r_dTRANSc + rasDiff*r_mTcT*(fRasCil-1) - rasDiff*(1-min(1,fRasCil))*mTcD

MuSC(0) = 100

TRANS(0) = 7600

DIFF(0) = 0

MuSCc(0) = 400

TRANSc(0) = 1900

```
********** MODEL PARAMETERS
mP = 0
mPTT = 0
mPcP = 0
mPcT = 0
mT = 0
mTDD = 0
mTcT = 0
mTcD = 0
d = 0
serumSwitch = 0
rasDiff=0
fRasCil=1

********** MODEL VARIABLES
serum = 1 - serumSwitch
sumPTD = MuSC + TRANS + DIFF + MuSCc + TRANSc
PAX7pos = MuSC + MuSCc
PAX7neg = TRANS + DIFF + TRANSc
pMuSC = (MuSC+MuSCc)/sumPTD * 100
pTRANS = (TRANS+TRANSc)/sumPTD * 100
pDIFF = DIFF/sumPTD * 100
pCil = (MuSCc+TRANSc)/sumPTD * 100

********** MODEL REACTIONS
r_mP = mP * MuSC *(1+serum)
r_mPTT = mPTT * MuSC *serumSwitch
r_mPcP = mPcP * MuSCc *serum
r_mPcT = mPcT * MuSCc *serumSwitch

r_mT = mT * TRANS *(1+serum)
r_mTDD = mTDD * TRANS *serumSwitch
r_mTcT = mTcT * TRANSc *serum
r_mTcD = mTcD * TRANSc *serumSwitch

r_dMuSC = d * MuSC
```

r_dMuSCc = d * MuSCc

r_dTRANS = d * TRANS

r_dTRANSc = d * TRANSc

r_dDIFF = d * DIFF

********** MODEL FUNCTIONS

********** MODEL EVENTS

********** MODEL MATLAB FUNCTIONS

### Original Data

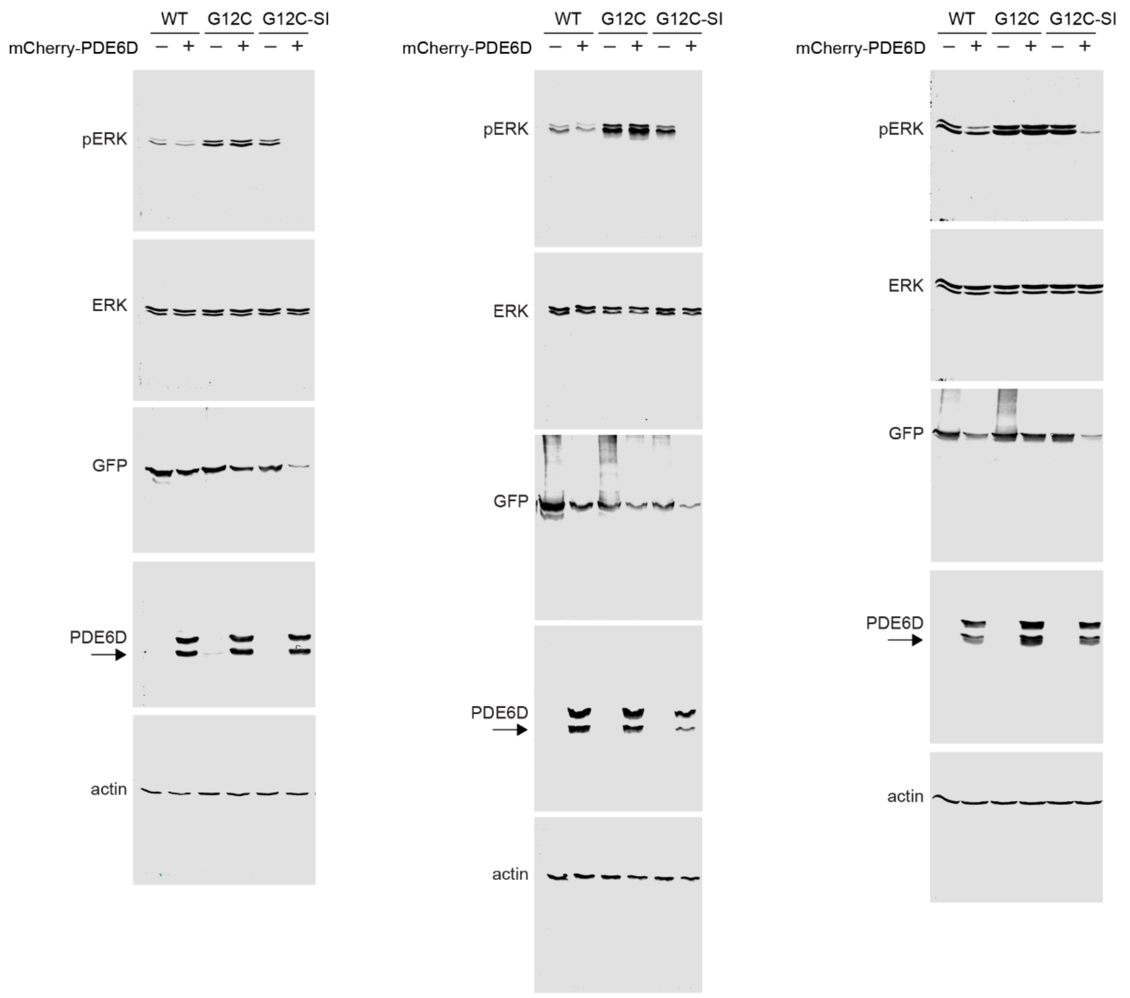


**Uncropped Western blot data. Related to Figure S5B.**

Whole blots of HEK cells transfected with EGFP-K-Ras (WT), EGFP-K-RasG12C (G12C) or EGFP-K-RasG12C-SI (G12C-SI) with or without co-transfection of mCherry-PDE6D at 1:1 DNA ratio for 24 h. Employed antibody labelings are indicated to the left of the blots. Arrows mark truncated/ cleaved protein consistently observed following transfection of mCherry-PDE6D.
